# Loss of region-specific glial homeostatic signature in prion diseases

**DOI:** 10.1101/823732

**Authors:** Natallia Makarava, Jennifer Chen-Yu Chang, Kara Molesworth, Ilia V. Baskakov

## Abstract

**Background:** Chronic neuroinflammation is recognized as a major neuropathological hallmark in a broad spectrum of neurodegenerative diseases including Alzheimer’s, Parkinson’s, Frontal Temporal Dementia, Amyotrophic Lateral Sclerosis, and prion diseases. Both microglia and astrocytes exhibit region-specific homeostatic transcriptional identities, which under chronic neurodegeneration, transform into reactive phenotypes in a region- and disease-specific manner. Little is known about region-specific identity of glia in prion diseases. The current study was designed to determine whether the region-specific homeostatic signature of glia changes with the progression of prion diseases, and whether these changes occur in a region-dependent or universal manner. Also of interest was whether different prion strains give rise to different reactive phenotypes.

**Methods:** To answer these questions, we analyzed gene expression in thalamus, cortex, hypothalamus and hippocampus of mice infected with 22L and ME7 prion strains using Nanostring Neuroinflammation panel at subclinical, early clinical and advanced stages of the disease.

**Results:** We found that at the preclinical stage of the disease, region-specific homeostatic identities were preserved. However, with the appearance of clinical signs, region-specific signatures were partially lost and replaced with a neuroinflammation signature. While the same sets of genes were activated by both prion strains, the timing of neuroinflammation and the degree of activation in different brain regions was strain-specific. Changes in astrocyte function scored at the top of activated pathways. Moreover, clustering analysis suggested that the astrocyte function pathway responded to prion infection prior to activated microglia or neuron and neurotransmission pathways.

**Conclusions:** The current work established neuroinflammation gene expression signature associated with prion diseases. Our results illustrate that with the disease progression, the region-specific homeostatic transcriptome signatures are replaced by region-independent neuroinflammation signature, which was common for prion strains with different cell tropism. The prion-associated neuroinflammation signature identified in the current study overlapped only partially with the microglia degenerative phenotype and the disease-associated microglia phenotype reported for animal models of other neurodegenerative diseases.

## Background

Chronic neuroinflammation is recognized as one of the major neuropathological hallmarks of neurodegenerative diseases including Alzheimer’s, Parkinson’s, Frontal Temporal Dementia, Amyotrophic Lateral Sclerosis, and prion diseases [1]. Chronic neuroinflammation manifests itself as a sustained activation of glial cells and the transformation of their homeostatic phenotype into reactive phenotypes [2, 3]. Transcriptome analysis and single-cell RNA-sequencing revealed considerable region-specific homeostatic heterogeneity in microglia and astrocyte phenotypes under normal conditions as well as dynamic phenotypic transformation in aging and neurodegenerative diseases [4–9]. While incredible progress has been made in characterizing diversity of glia phenotypes using mouse models of neurodegenerative diseases, concerns whether mouse models faithfully recapitulate key aspects of disease in human have been raised on numerous occasions [10–12].

For elucidating mechanisms behind chronic neuroinflammation and neurodegeneration, prion disease offers several advantages over other neurodegenerative disorders. The most obvious reason behind choosing prion disease is that animals inoculated with prions develop actual *bona fide* prion disease, not a disease model [13]. Inbred mice infected with prions recapitulate neuropathological and biochemical features associated with naturally occurring prion diseases including prion diseases of humans. Prion diseases can be efficiently transmitted between wild type animals or inbred laboratory mice, the process that does not rely on expression or overexpression of modified human genes.

Prions, or PrP^Sc^, are proteinaceous infectious agents that consist of misfolded, self-replicating states of a sialoglycoprotein called the prion protein or PrP^C^ [14, 15]. Prion diseases display diverse disease phenotypes, a feature attributed to the ability of PrP^C^ to acquire multiple, conformationally distinct, self-replicating PrP^Sc^ states referred to as prion strains or subtypes [16–18]. In addition to differences in structure, prion or PrP^Sc^ strains exhibit different patterns of terminal carbohydrate groups and a variable density of sialic acid residues on their surface [19–22]. The differences in surface-exposed carbohydrate epitopes are believed to be due to a selective strain-specific recruitment of PrP^C^ molecules among a large pool of more than 400 PrP^C^ sialoglycoforms expressed by a cell [19, 21, 23]. Considering the structural diversity of PrP^Sc^ and the diversity of carbohydrate epitopes on a surface of PrP^Sc^ particles, it is not surprising that prion strains exhibit selective strain-specific tropism with respect to brain region and cell type [24–26].

Previous studies analyzed transcriptome changes using laboratory inbred mice infected with prions [27–33]. Analyses of whole transcriptome, in combination with analyses of selective gene sets, identified activation of microglia with strong proinflammatory characteristics as a common signature of chronic neuroinflammation associated with prion diseases [24, 29, 30, 32, 34–36]. While these studies revealed a variety of differentially activated genes and pathways, our knowledge about reactive phenotypes of microglia and astrocytes in prion diseases is very limited in comparison to other neurodegenerative diseases. In the majority of previous studies, whole brain tissues were used for transcriptome analysis leaving region specific identities concealed. In the current study, we asked whether prion strains give rise to different reactive phenotypes in glia and whether these phenotypes are influenced by region-specific homeostatic signatures. To address these questions, we analyzed gene expression in four brain regions in mice infected with two prion strains, 22L and ME7, that have different cell- and region-specific tropism [24], using a Nanostring Neuroinflammation panel. At the preclinical stage of the disease, the region-specific homeostatic identity of glia was well preserved. However, with the disease progression, the region-specific homeostatic signatures were partially lost and replaced with neuroinflammation signature. The same genes were activated by both prion strains, however, the timing of neuroinflammation and the degree of activation in different brain regions were strain-specific. Global significance scoring of differentially expressed genes identified Astrocyte Function pathway at the top of the list followed by Inflammatory Signaling, Matrix Remodeling, and Activated Microglia. Moreover, clustering analysis of gene expression patterns suggested that Astrocyte Function pathway responded to prion infection prior to Activated Microglia or Neuron and Neurotransmission pathways. The current work established Neuroinflammation gene expression signature associated with prion diseases and demonstrated that it was independent of brain region or prion cell tropism.

## Methods and Materials

### Animal experiments and brain tissue collection

Using isoflurane anesthesia, six-week-old C57BL/6J female mice were inoculated intraperitoneally with 200 μl of 1% 22L or ME7 brain homogenate in PBS, pH 7.4. A control group was inoculated with PBS only. Animals were regularly scored for signs of neurological impairment and disease progression: progressive difficulty walking on a beam, hind limb clasping, and weight loss. Pre-symptomatic 22L and ME7 samples were collected at 153 – 154 days post-inoculation (dpi) from animals showing no clinical signs and no weight loss. The early clinical 22L samples were collected upon consistent observation of mild motor impairment signs for two weeks, which were the first clinical signs observed. No significant weight loss was observed for these animals. For ME7, 2^nd^ time point samples were collected at 224 dpi from mice displaying no signs of neurological impairment or weight loss. ME7 mice started to develop first clinical signs of the disease at 280 – 343 dpi. Within 15 – 31 days after first clinical signs, 22L and ME7 mice became unable to walk on a beam, developed abnormal gait and became lethargic. Mice were considered terminally ill when they were unable to rear and lost 20% of their weight. At this time, 3^rd^ point samples were collected. Mice were euthanized by CO_2_ asphyxia and decapitation.

After euthanasia, brains were immediately extracted and kept ice-cold during dissection. Brains were sliced using rodent brain slicer matrix (Zivic Instruments, Pittsburg, PA). 2 mm central coronal sections of the brain were used to collect individual regions. Allen Brain Atlas digital portal (http://mouse.brain-map.org/static/atlas) was used as a reference. Hypothalami (HTh), as well as left and right thalami (Th), hippocampi (Hp) and cortexes (Ctx) were collected into RNase-free sterile tubes, frozen in liquid nitrogen, and stored at −80°C until RNA isolation. The anterior part of the brain and left half of the posterior part of the brain were saved in 10% buffered formalin for immunohistochemistry. Dissection remnants were frozen in a separate tube for Western blot with anti-PrP antibody ab3531 (Abcam, Cambridge, MA)

### RNA isolation

Brain tissue samples were homogenized within RNase-free 1.5 ml tubes in 200 µl of Trizol (Thermo Fisher Scientific, Waltham, MA, USA), using RNase-free disposable pestles (Fisher scientific, Hampton, NH). After homogenization, an additional 600 µl of Trizol was added to each homogenate, and the samples were centrifuged at 11,400 × g for 5 min at 4°C. The supernatant was collected, incubated for 5 min at room temperature, then supplemented with 160 µl of cold chloroform and vigorously shaken for 30 sec by hand. After additional 5 min incubation at room temperature, the samples were centrifuged at 11,400 × g for 15 min at 4°C. The top layer was transferred to new RNase-free tubes and mixed with an equal amount of 70% ethanol. Subsequent steps were performed using Aurum Total RNA Mini Kit (Bio-Rad, Hercules, CA, USA) following manufacturer’s instructions. Isolated total RNA was subjected to DNase I digestion. RNA purity and concentrations were estimated using NanoDrop One Spectrophotometer (Thermo Fisher Scientific, Waltham, MA, USA).

### NanoString

Samples were processed by the Institute for Genome Center at the University of Maryland School of Medicine using nCounter Mouse Neuroinflammation Panel. Only samples with RNA integrity number RIN > 7.2 were used for Nanostring. All data passed QC, with no imaging, binding, positive control, or CodeSet content normalization flags. Analysis of data was performed using nSolver Analysis Software 4.0, including nCounter Advanced Analysis (version 2.0.115). For agglomerative clusters and heat maps, genes with less than 10% of samples above 20 counts were excluded. Z-score transformation was performed for genes. Clustering was done using Euclidian distance, linkage method was average.

### Histopathological study

Formalin-fixed brain sections were submerged for 1 hour in 95% formic acid to deactivate prion infectivity before being embedded in paraffin. Subsequent 4 µm sections produced using Leica RM2235 microtome were mounted on slides and processed for immunohistochemistry. To expose epitopes, slides were subjected to 20 min hydrated autoclaving at 121°C in trisodium citrate buffer, pH 6.0, with 0.05% Tween 20. Rabbit anti-Iba1 (Wako, Richmond, VA) was used to stain microglia. Chicken polyclonal anti-GFAP (Sigma-Aldrich, St. Louis, MO) was used to stain astrocytes. For detection of disease-associated PrP, 5 min treatment with 88% formic acid was used following autoclaving. PrP was stained with anti-prion antibody SAF-84 (Cayman Chemical, Ann Arbor, MI). Detection was performed using DAB Quanto chromogen and substrate (VWR, Radnor, PA).

## Results

### Experimental design

C57Black/6J mice were intraperitoneally (IP) inoculated with 22L or ME7 mouse-adapted prion strains (200 μl, 1% brain homogenate) at 5 weeks old and euthanized at three time points post inoculation (Fig. S1A, Table S1). Animals infected with 22L prions were euthanized at the preclinical (1^st^ time-point, 153 days post-inoculation (dpi)), early clinical (2^nd^ time-point, 186 - 197 dpi) and advanced clinical stages of the disease (3^rd^ time-point, 168 – 225 dpi) (Table S1). ME7-infected animals were euthanized at the early preclinical (1^st^ time-point, 154 dpi), late preclinical (2^nd^ time-point, 224 dpi) and advanced clinical stages of the disease (3^rd^ time-point, 295 - 363 dpi) (Table S1). For identifying early stages, advanced clinical stages and monitoring progression of the disease, the disease scoring protocol was employed as described in Methods. IP inoculation allowed us to avoid effects related to brain trauma, yet this route of infection had some drawbacks such as differences in the onset of the disease and relatively poor cooperativity in disease progression within an animal group. In animals inoculated with 22L prions, the timing of the early and advanced clinical stages overlapped between groups due to variations of the disease onset and the rate of the disease progression (Fig. S1A, Table S1). As control groups, C57Black/6J mice were inoculated IP with PBS and euthanized at 151 dpi (controls for the 1^st^ time-point for 22L and ME7), 197 - 223 dpi (controls for the 2^nd^ time-point for 22L and ME7, and for the 3^rd^ time-point for 22L) and 295 - 363 dpi (controls for the 3^rd^ time-point for ME7) (Fig. S1A, Table S1).

For assessing region-specific neuroinflammation status, four brain regions - thalamus (Th), hypothalamus (HTh), cortex (Ctx) and hippocampus (Hp) (n=3 individual animals per group) were selected based on previous studies [24, 25, 37, 38] (Fig. S1B). Analysis of the expression of genes associated with neuroinflammation was conducted using nCounter Nanostring Neuroinflammation panel (Table S2) that analyzes expression of 757 genes (including 13 housekeeping genes), which assess 23 pathways including Activated Microglia, Innate Immune Response, Adaptive Immune Response, Growth Factor Signaling, Inflammatory Signaling, Apoptosis, Autophagy, Astrocyte Function and others (the full list is in Fig. S2).

### At the preclinical stage, the neuroinflammation gene expression profile displays region-specific identity

Agglomerative hierarchical clustering of all data collected at the first, preclinical time point reveled that four brain regions displayed region-specific gene expression profiles, illustrating homeostatic signatures of individual regions (Fig. 1). Four distinctive clusters corresponding to Th, HTh, Ctx and Hp were observed (Fig. 1). With an exception of the Th cluster, 22L and ME7 datasets did not segregate into separate sub-clusters in remaining regions, but were mixed with normal controls. This result suggests that the thalamus might be the first region affected by neuroinflammation. A small subset of genes covered by the panel showed minor yet statistically significant up- or down-regulation at preclinical stages (Table S3). However, these changes were not sufficient to override region-specific homeostatic identity. In summary, at the preclinical stage, the region-specific homeostatic identities were well-preserved in all brain regions (Fig. 1). The largest proportion of genes covered by the Neuroinflammation panel report on microglia phenotype and their activation state (Fig. S2). As such, the region-specific homeostatic signatures are indicative of differences in microglia phenotypes in the four brain regions (Fig. 1).

**Figure 1.**
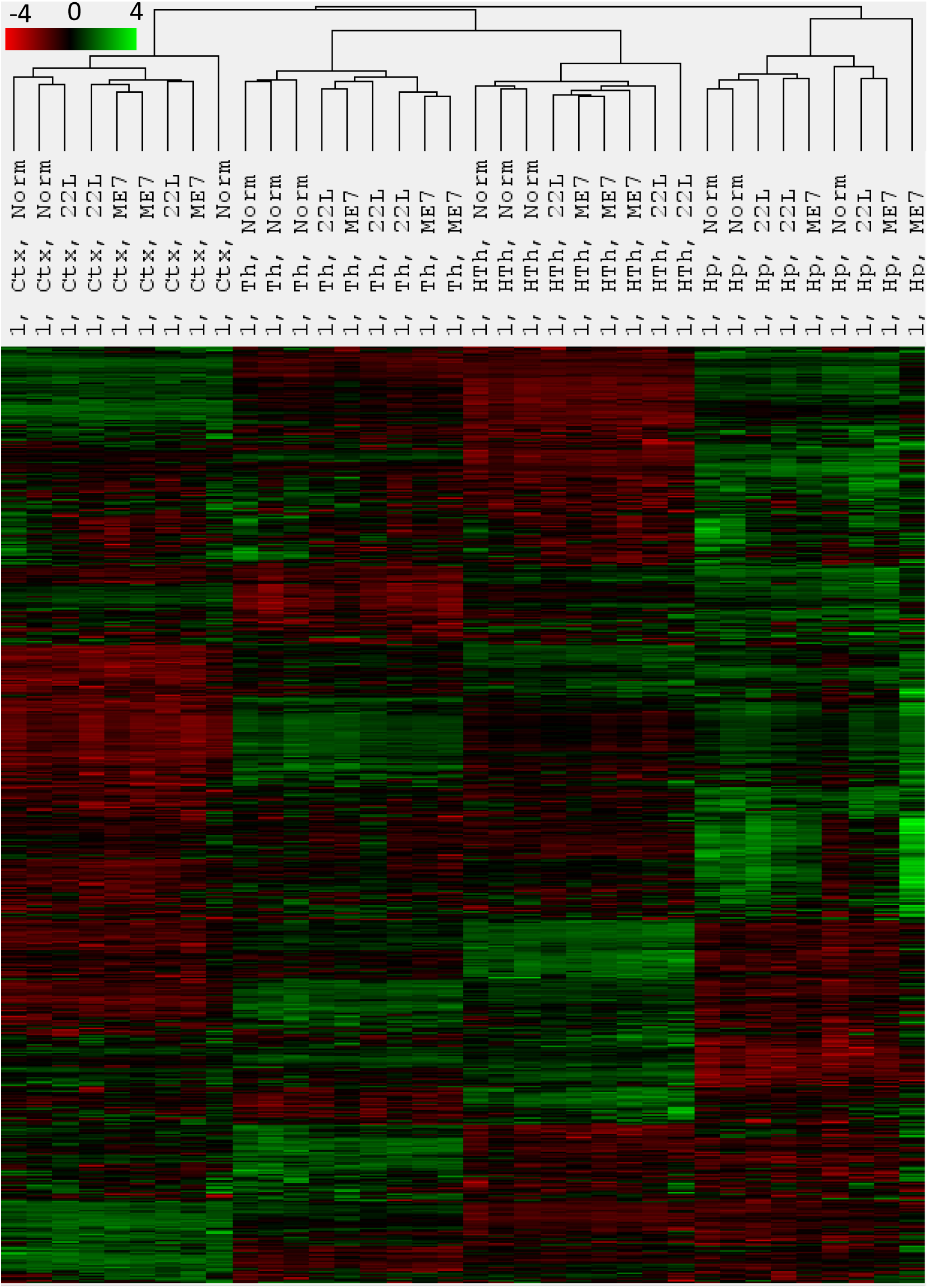
Agglomerative hierarchical cluster analysis of all data collected for the first, preclinical time point. Four well-defined clusters formed strictly according to the brain region presenting region-specific homeostatic signatures. Within each region-specific cluster, 22L- and ME7-infected animals did not cluster away from the age-matched control (Norm) animals with the exception of thalamus.

### Gradual loss of the region-specific homeostatic signatures with the disease progression

Agglomerative hierarchical clustering of all data collected at the second time point revealed a group of upregulated genes (Fig. S3, orange frame). Th and HTh from one 22L-infected animal (animal #4) clustered separately from all other samples forming a well-separated branch (Fig. S3, red shading). Within its group, the animal #4 showed the highest amounts of PrP^Sc^ on Western blot (Fig. S1C) and the most pronounced inflammation of microglia as assessed by immunostaining of brain sections using microglia-specific marker Iba1 (Fig. S4). Ctx of this animal also showed upregulation of the same set of genes, although to a lower degree (Fig. S3). Upregulation of the same genes were also visible in the thalamus of 22L-infected animal #5, although this sample remains in the cluster with other thalami. Notably, the ranking order between individual animals of the 22L group was the same (the most affected #4>#5>#6) regardless of whether it was assessed by differential gene expression, the amounts of PrP^Sc^ by Western blot, or microglia activation by immunostaining with Iba1 (Fig. S1B, S3, S4). In ME7-infected animals, which were all asymptomatic at the second time point, Th also showed upregulation of the same set of genes as 22L-affected Th, with one ME7-affected Th (animal #13) clustering with 22L-infected Th (animal #5) (brown shading in Fig. S3). To summarize, at the time when the signs of the disease began to appear, expression of a subset of genes started to dominate over the region-specific homeostatic signature in animals that were most advanced in the disease progression. The same set of genes appears to be upregulated regardless of the brain region or prion strain.

The trend noticed at the second time point strengthened further at the third time point. At the advanced stage, 22L Th and HTh from all three animals of the group, now joined by the three ME7 Th of its group, clustered away from all other samples (Fig. 2, red shading). Notably, the upregulation of the gene set defined by a red frame was so profound that it overrode region-specific transcriptome signatures still recognizable in the 22L Th+HTh and ME7 Th sample sets (Fig. 2). The set of genes that drove separation of 22L Th+HTh and ME7 Th into a distant cluster will be referred to as neuroinflammation signature associated with prion disease (marked by red frame in Fig. 2). Ctx of all 22L- and ME7-infected animals also showed upregulation of the genes within the neuroinflammatory signature block, yet to a lower degree, and formed a sub-cluster within the Ctx cluster (Fig. 2). These results suggest that at the advanced stage of the diseases, in the brain regions that are the most strongly affected by prions, glia partially lose their region-specific homeostatic signature and merge into a highly reactive phenotype. This reactive phenotype is characterized by upregulation of the same set of genes as defined by the neuroinflammatory signature block. The full list of differentially expressed genes is presented in Table S4. The genes within the neuroinflammation signature block were common for all regions, although the extent of upregulation varied in a region-specific manner as discussed below.

**Figure 2.**
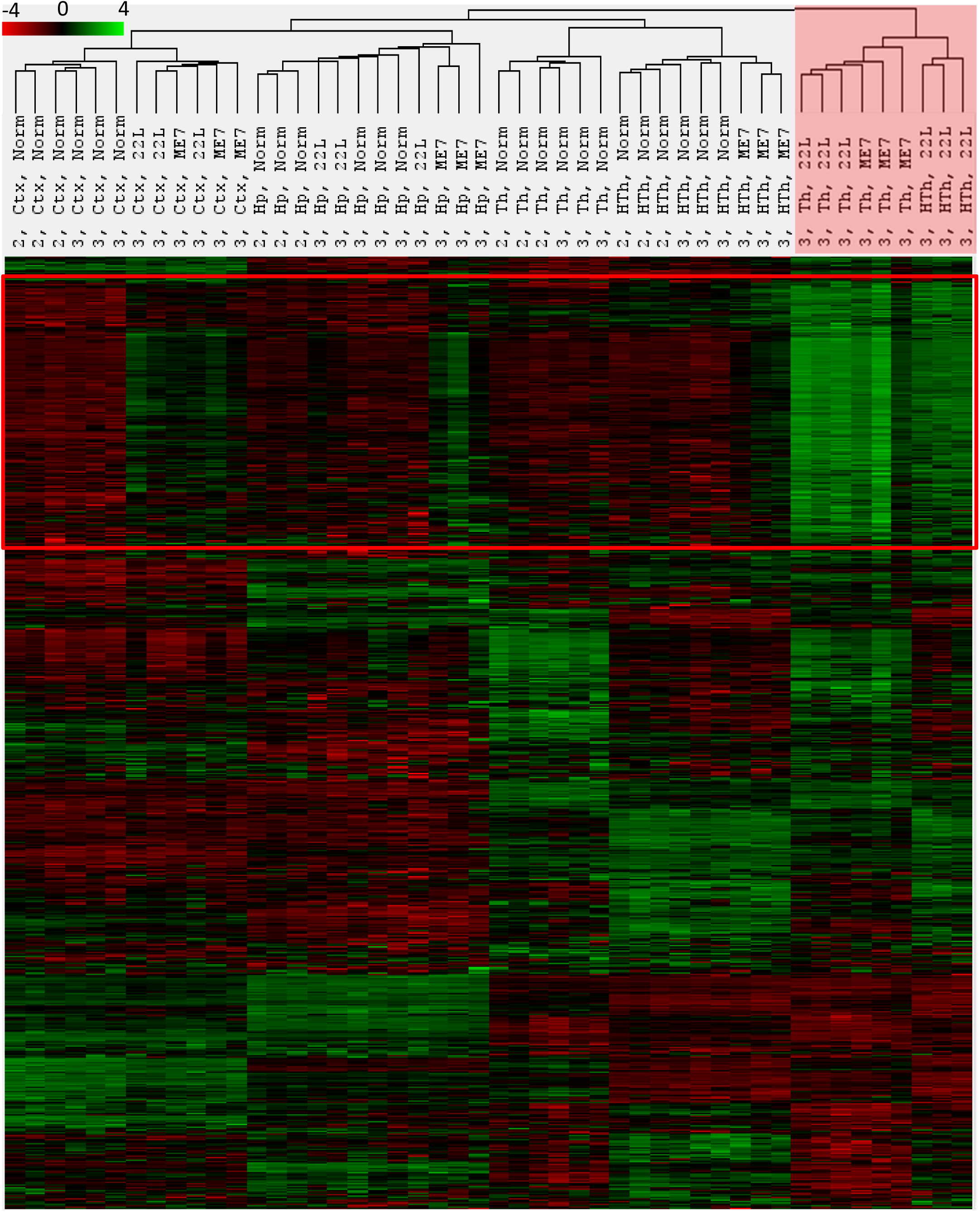
Agglomerative hierarchical cluster analysis of all data collected for the third time point. The red frame define genes forming the neuroinflammatory signature. 22L Th, 22L HTh and ME7 Th clustered away from all other samples as indicated by red shading. This separation is driven by a strong upregulation of genes within the neuroinflammatory signature block. Notably, 22L Ctx and ME7 Ctx, Hp and HTh also show upregulation of the same genes, which separates these samples from normal controls within the corresponding regional clusters.

### Region- and strain-specific dynamics of neuroinflammation

Agglomerative clustering of grouped samples (n=3 per group) collected for four brain regions in both strains at three time points showed the same dynamics as clustering of individual samples (Fig. 3). Again, clear separation of 22L Th, 22L HTh and ME7 Th into a highly distinctive cluster was evident in samples from the advanced stage of the disease (Fig. 3). Notably, Ctx, Hp and HTh showed upregulation of the same sets of genes as those found in Th, although to a lesser degree than in Th, and to a different extent in 22L compared to ME7 (Fig. 3).

**Figure 3.**
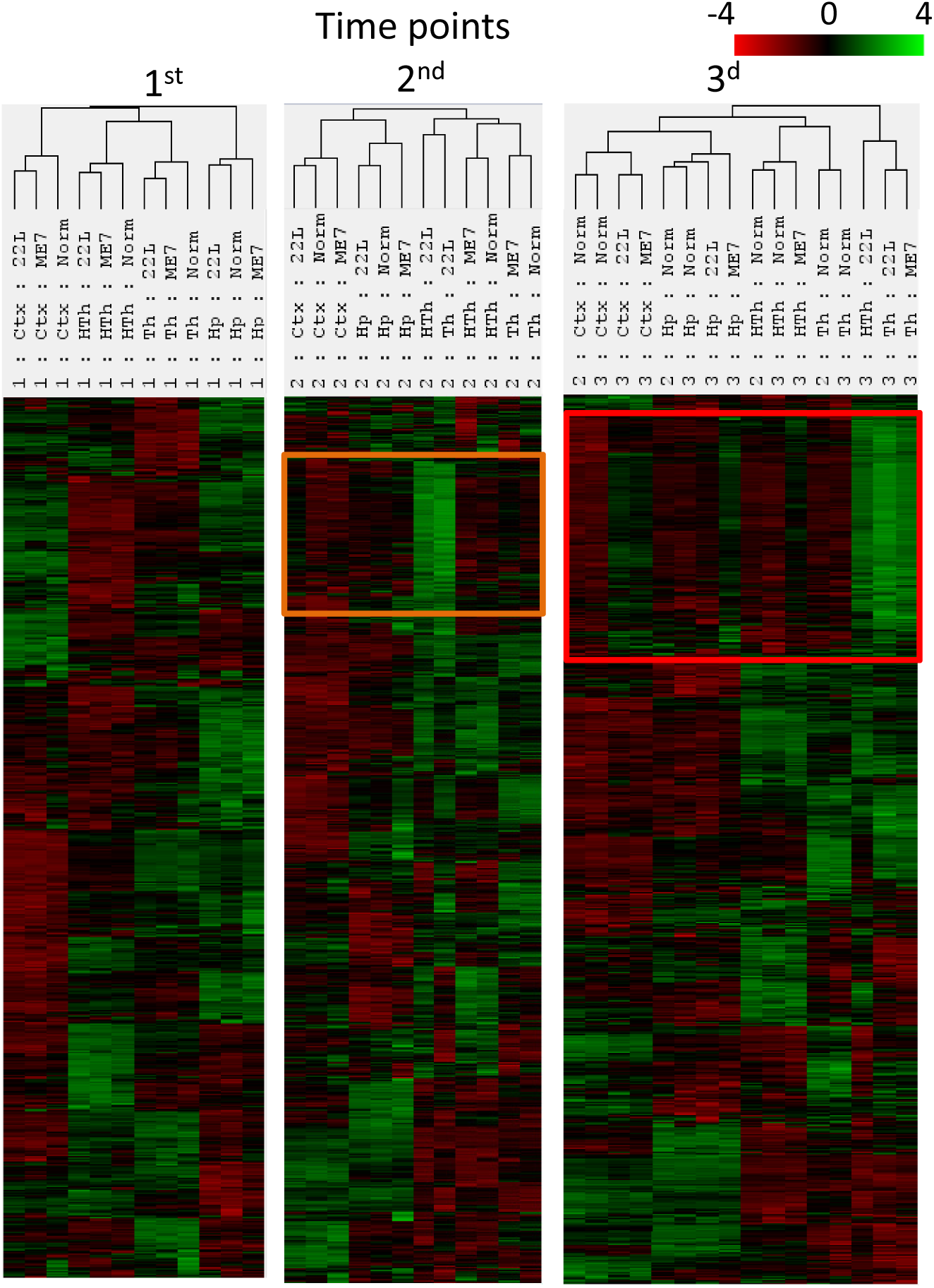
Agglomerative clustering analysis of grouped samples collected for all three time points. To group samples, the geometric mean of expression levels for all samples from each group (n=3) were calculated. The set of genes upregulated at the second time point (orange frame) expands further at third time point (red frame).

Analysis of genes within the neuroinflammation signature block revealed that the majority of genes belonged to the Astrocyte Function, Microglia Activation, Inflammatory Signaling and Autophagy pathways (Fig. 4). The genes within the neuroinflammation signature block responded to prion infection in a coherent manner and showed the same dynamics, as assessed across four brain regions, three time points and two prion strains (Fig. 4).

**Figure 4.**
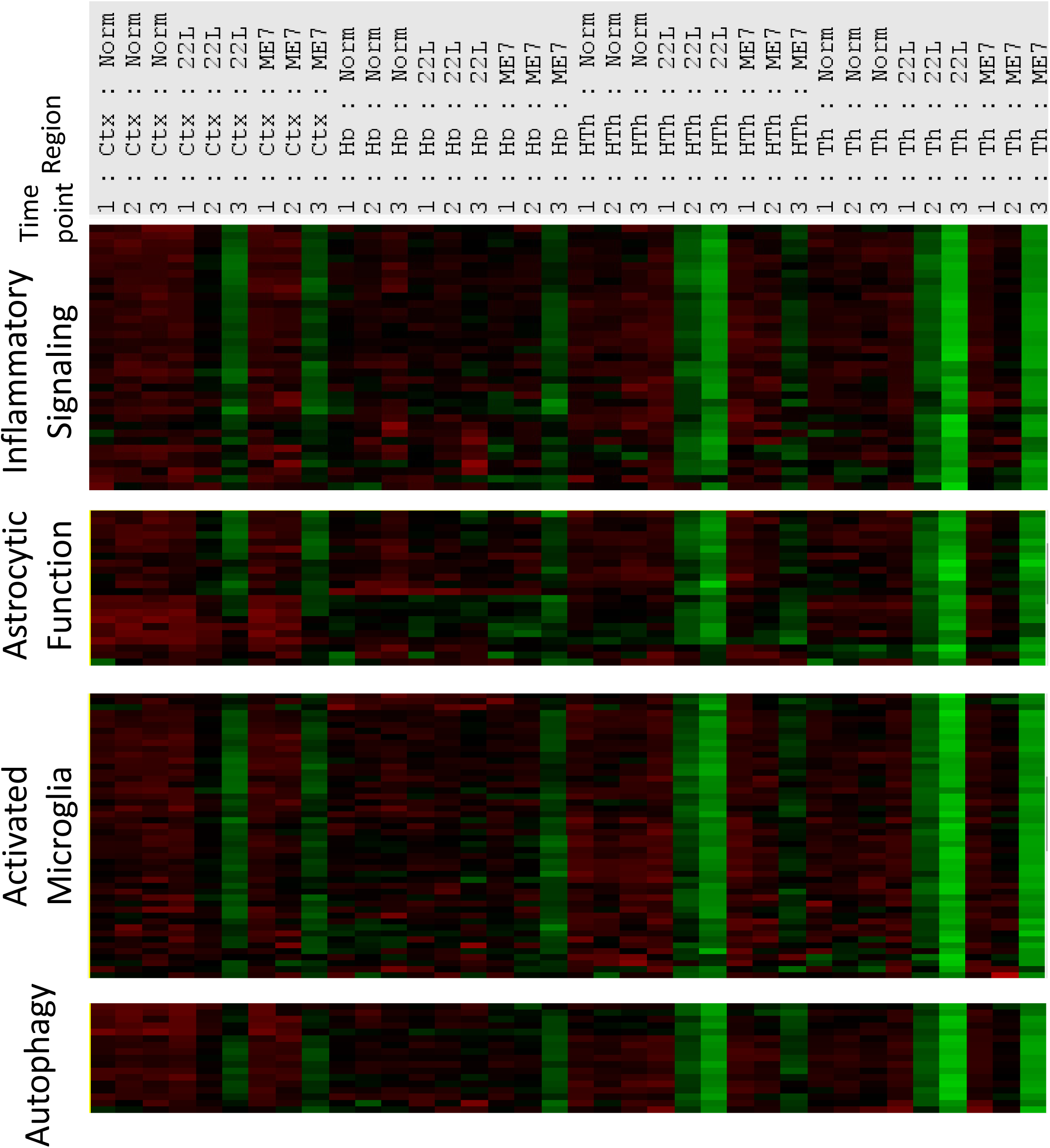
Gene composition within the neuroinflammation signature block. Distribution of upregulated genes within the neuroinflammation signature block according to pathways. Grouped samples corresponding to four brain regions in animals infected with 22L or ME7 strains or non-infected controls (Norm) and collected at three time points are presented.

Close comparison of 22L and ME7 sets revealed that the same genes were activated by both strains. However, the timing of neuroinflammation and the degree of activation in different brain regions were strain-specific. To establish a strain-specific ranking order with respect to (i) the temporal spread of neuroinflammation across the brain and (ii) the extent to which brain regions were affected at the advanced stage, we first assessed the intensity of changes within the genes of the neuroinflammation signature block. For 22L, the ranking order was Th>HTh>Ctx>>Hp (where Th was the earliest and most affected region), whereas for ME7, the ranking order was Th>Ctx>Hp=HTh (Fig 5A). The ranking order established by the gene expression correlated well with the region-specific deposition of PrP^Sc^, reactive microgliosis and astroglyosis, as assessed by staining for Iba1 and GFAP, respectively (Fig. S5). Counting a number of differentially expressed genes at the advanced stage of disease (P< 0.05, fold change > +/-1.2) showed the same ranking order as assessed by intensity of differential gene expression (Fig 5B). This ranking order was also confirmed upon examination of region-specific expression of individual genes including *Cxcl10, Serpina3n, Cd68, Clec7a* (Fig. S6).

**Figure 5.**
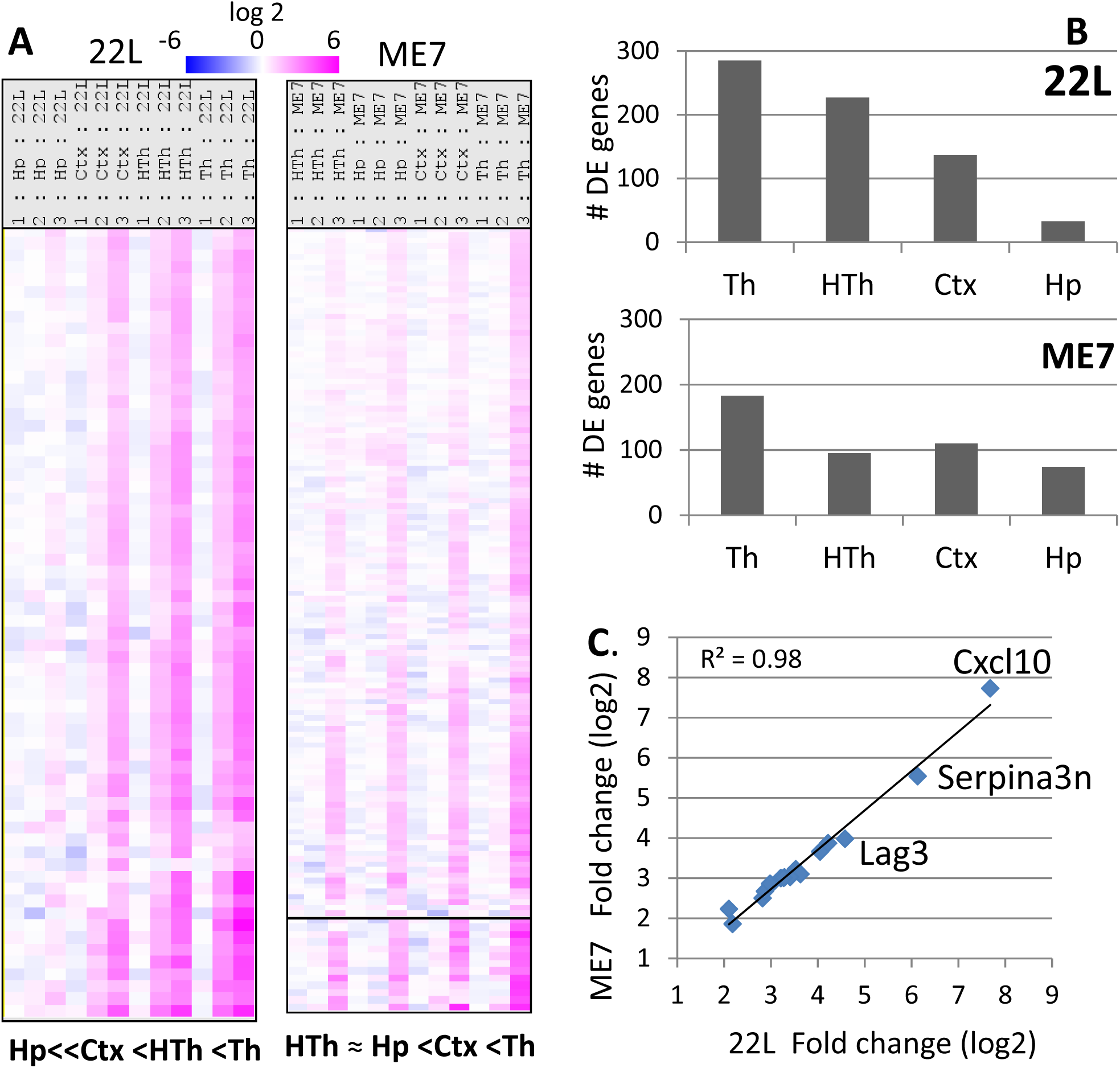
Region- and strain-specific dynamics of neuroinflammation. **(A)** Strain-specific ranking order with respect to the temporal spread of neuroinflammation across four brain areas and the degree to which the brain regions were affected at the advanced stage of the disease, as judged by the sum intensity of changes. The data presented as heatmap of grouped samples (n=3) showing log2 fold change in 22L- or ME7-infected mice over normal controls. **(B)** Number of differentially expressed (DE) genes in four brain regions in 22L- and ME7-infected mice at the advanced stage of the disease. **(C)** Correlation between log2 fold change of gene expression between ME7- and 22L-infected animals for the top 20 differentially expressed genes at the advance stage of the disease.

Analysis of top activated genes in 22L Th and ME7 Th revealed excellent correlation between the two strains with R^2^ = 0.98 (Fig. 5C, Table 1). Moreover, 22L and ME7 did not separate into different sub-clusters illustrating lack of strain-specificity in the overall pattern of gene activation (Fig. 2). For instance, 22L Ctx and ME7 Ctx displayed similar pattern of gene activation and together formed a sub-cluster within the Ctx cluster (Fig. 2). To summarize, these data strongly indicate lack of strain-specificity in neuroinflammatory response. The strain-specificity consisted of the differences in tropism to different brain regions and the degree of gene activation rather than an activation of different subsets of genes.

**Table 1.**
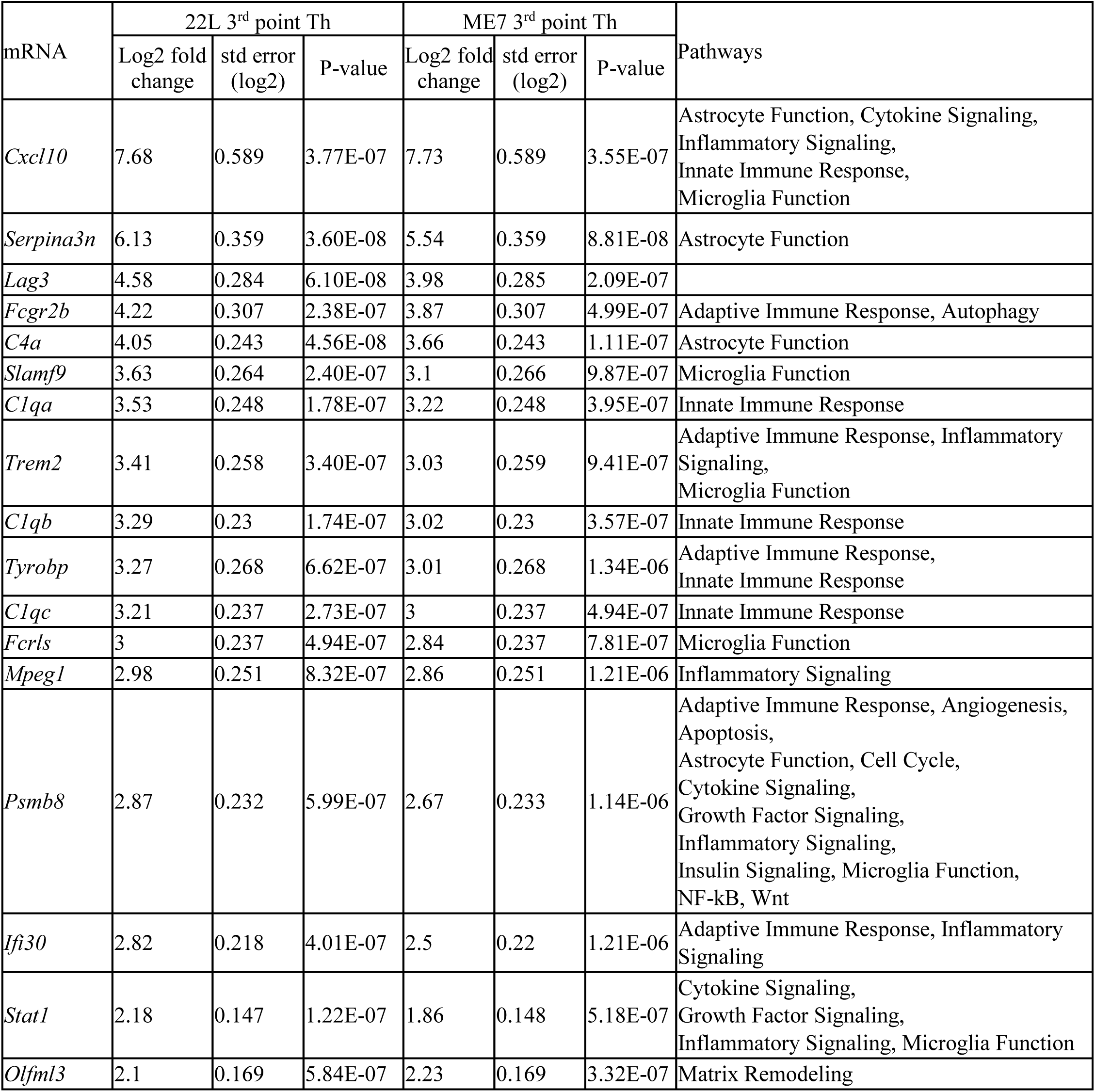
Top differentially expressed genes in 22L- and ME7-infected animals at the advanced stage of the disease.

### Change in astrocyte function scores at the top of the pathways analyzed

Among the pathways covered by the Neuroinflammation panel, we wanted to know what pathways were affected the most. To answer this question, we focused on the thalamus, which showed the strongest activation among four brain regions analyzed. The heatmap of pathway scores revealed that three pathways (Neuron and Neurotransmission, Epigenetic regulation, and Oligodendrocyte function) were downregulated, whereas the remaining pathways were strongly upregulated in both 22L and ME7 relative to the controls (Fig. 6). Undirecteded global significance scores of differences between prion-infected and control animals identified Astrocyte Function pathway at the top of the list followed by Inflammatory Signaling, Matrix Remodeling, Adaptive Immune Response and Microglia Function pathways for both prion strains (Table 2). Innate Immune Response, NF-kB and Autophagy pathways also scored highly. The undirected global significance scores of the genes related to Inflammatory Signaling, Astrocyte Functions and Activated Microglia were much higher than the scores for the Neurons and Neurotransmission pathway that was close to the bottom of the list (Table 2).

**Figure 6.**
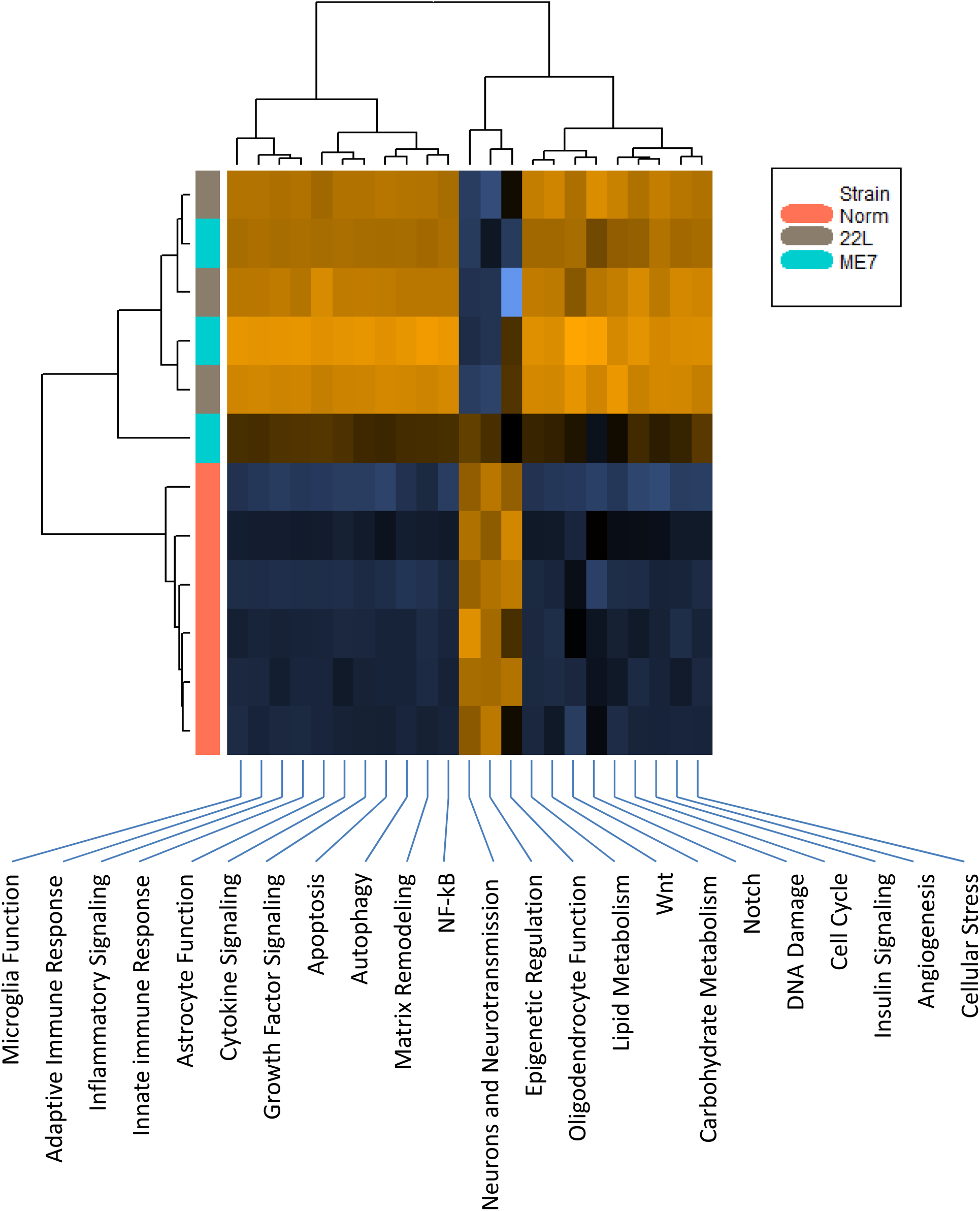
Heatmap of the pathway scores. The pathways were scored based on the datasets generated for thalamus at the third time point. Three pathways (Neurons and Neurotransmission, Epigenetic Regulation, and Oligodendrocyte Function) were downregulated, whereas remaining pathways were upregulated in 22L- and ME7-infected animals. Upregulated pathways are shaded brown, downregulated pathways are shaded blue.

**Table 2.** Undirected and directed Global Significance Scores of 22 pathways analyzed by the Neuroinflammation panel.

| Gene sets | Undirected GSS * |  | Directed GSS * |  |
| --- | --- | --- | --- | --- |
|  | 22L vs. Norm | ME7 vs. Norm | 22L vs. Norm | ME7 vs. Norm |
| Astrocyte Function | 6.42 | 5.626 | 5.875 | 5.558 |
| Inflammatory Signaling | 6.003 | 5.455 | 5.891 | 5.429 |
| Matrix Remodeling | 5.689 | 5.091 | 5.241 | 4.942 |
| Adaptive Immune Response | 5.512 | 4.976 | 4.861 | 4.666 |
| Autophagy | 5.06 | 4.422 | 4.388 | 4.04 |
| Microglia Function | 5.055 | 4.465 | 4.729 | 4.234 |
| Innate Immune Response | 5.035 | 4.491 | 4.582 | 4.314 |
| Cytokine Signaling | 4.778 | 4.274 | 4.241 | 4.026 |
| NF-kB | 4.684 | 4.275 | 4.529 | 4.236 |
| Insulin Signaling | 4.557 | 3.863 | 3.301 | 3.272 |
| Angiogenesis | 4.556 | 3.851 | 3.305 | 3.168 |
| Wnt | 4.363 | 3.256 | 1.396 | 2.421 |
| Growth Factor Signaling | 4.327 | 3.733 | 3.426 | 3.289 |
| Lipid Metabolism | 3.916 | 3.146 | 2.803 | 2.625 |
| Apoptosis | 3.469 | 2.879 | 2.074 | 2.327 |
| Cellular Stress | 3.22 | 2.678 | 1.889 | 1.818 |
| Neurons and Neurotransmission | 3.002 | 2.503 | -1.052 | 1.241 |
| Cell Cycle | 2.997 | 2.368 | 2.019 | 2.002 |
| Notch | 2.73 | 2.053 | 1.786 | 1.764 |
| Carbohydrate Metabolism | 2.655 | 2.402 | 1.597 | 2.259 |
| DNA Damage | 2.162 | 1.639 | 1.304 | 1.325 |
| Epigenetic Regulation | 2.116 | 1.469 | -0.968 | 0.834 |

In prion diseases, chronic inflammation is accompanied by proliferation of microglia [39]. To test whether the global significance scores of top pathways reflect possible changes in cell type composition in addition to differential gene expression per se, next we compared cell population scores of microglia, astrocytes and neurons (Fig. 7). In 22L- and ME7-infected animals, microglial cell specific markers scored significantly higher relative to the controls. This result is consistent with significant proliferation and/or infiltration of microglia. As a result of proliferation/infiltration, the specific gene set scores for the Activated Microglia pathway and other pathways dominated by microglia-specific genes might have been inflated (Fig. 7). Despite a substantial increase in the specific gene set score for the Astrocyte Function pathway, an increase in cell population score for astrocytes in infected relative to control animals was considerably less profound in comparison to those of microglia (Fig. 7). Unlike microglia, astrocytes do not proliferate in prion diseases [40]. A modest increase in scoring of the astrocyte markers is likely to be attributed to astrocyte hypertrophy. A global score of neuron-specific markers did not show a significant drop, yet notable downregulation of the Neurons and Neurotransmission pathway was detected in prion-infected animals (Fig. 7). However, close examination of the fold change in expression of individual genes associated with the Neurons and Neurotransmission pathway was mostly small and of low statistical significance (Fig. S7). Few genes in the Neurons and Neurotransmission pathway showed strong upregulation. These genes also belong to other pathways and their activation was most probably related to the upregulation of these other pathways (*C3ar1* and *P2rx7* – activated microglia, *S1pr3* – astrocyte function, etc., Fig.S7).

**Figure 7.**
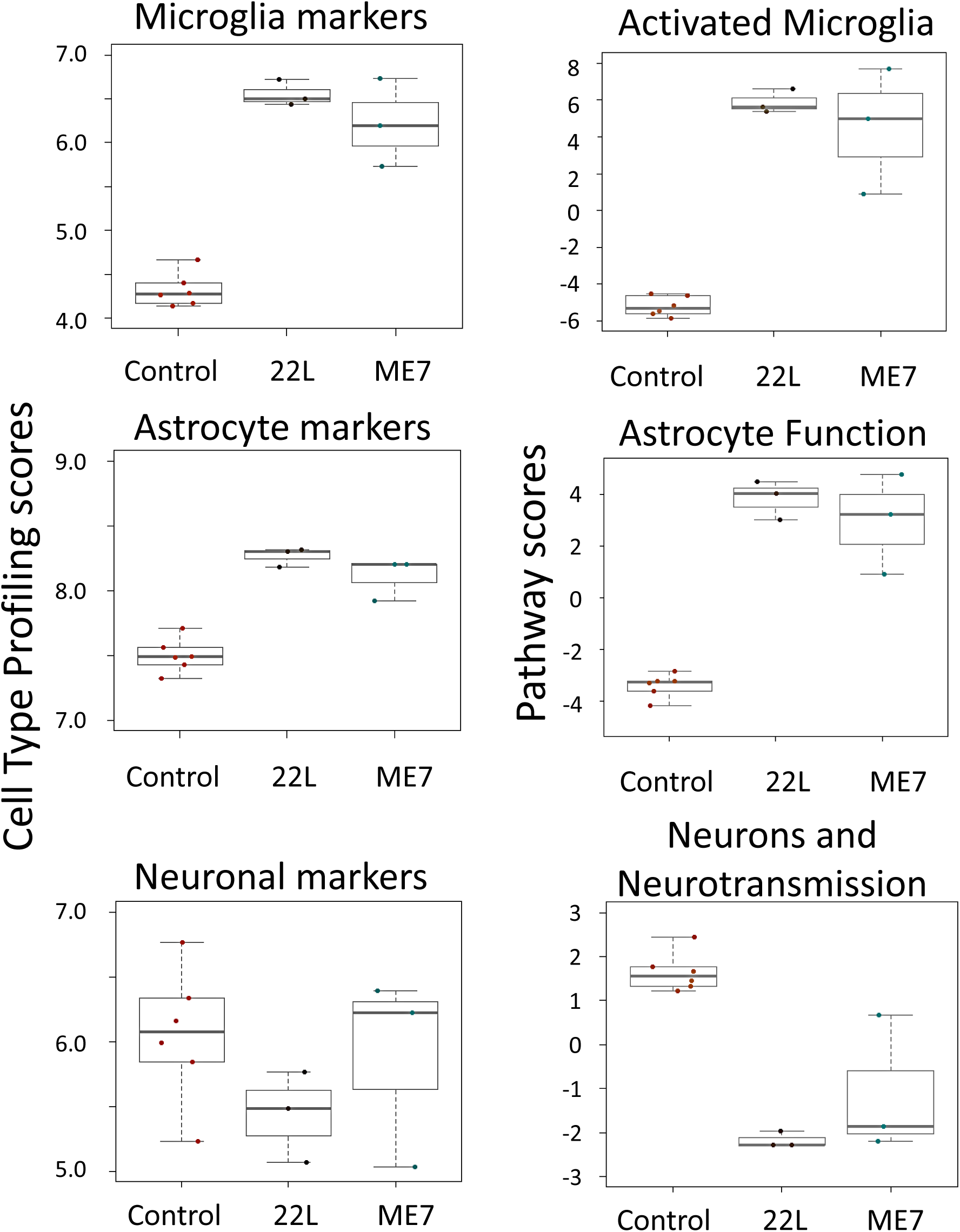
Cell population scores and pathway scores. Cell population scores for microglia, astrocytes and neurons, as well as scores for Activated Microglia, Astrocyte Function and Neuron and Neurotransmission pathways were generated based on the datasets taken for thalamus at the third time point.

To find what cell types (microglia, astrocytes or neurons) respond to prion infection at the preclinical stage of the disease, nCounter Advanced Analysis was employed to cluster samples based on gene expression pattern in Astrocyte Function, Microglia Function and Neurons and Neurotransmission pathways (Fig. 8). Clustering based on the genes in the Astrocyte Function pathways revealed that thalami from normal and disease-affected animals separated into two clusters already at the first time point and continued to cluster away from each other for the second and third time points (Fig. 8). In contrast, when genes associated with Microglia Function or Neurons and Neurotransmission were assessed, heatmaps demonstrated that normal and prion-affected thalami clustered away from each other only at the third, advanced stage of the disease (Fig. 8).

**Figure 8.**
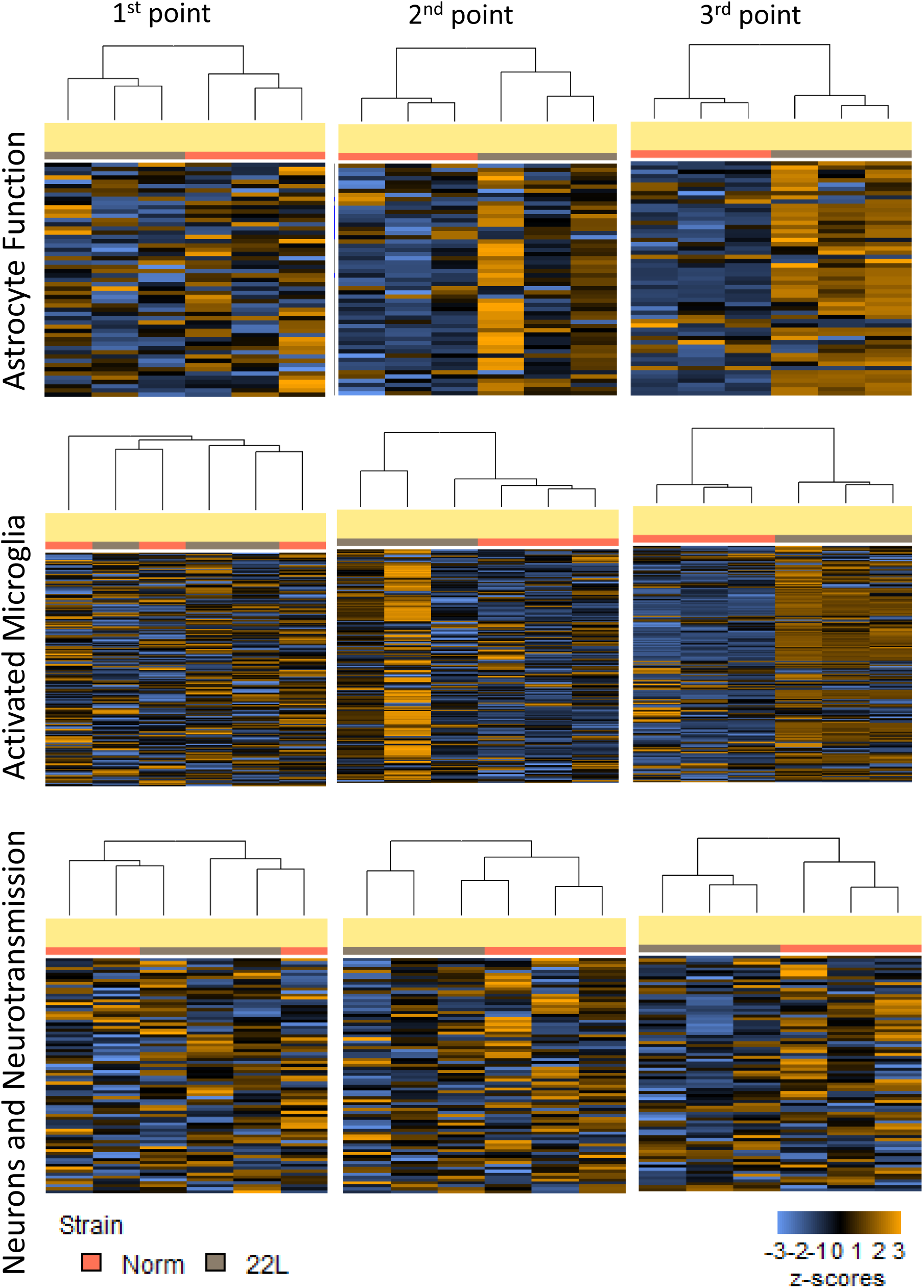
Hierarchical cluster analysis of the pathways. Hierarchical cluster analysis of the Astrocyte Function, Activated Microglia and Neuron and Neurotransmission pathways was performed based on the datasets generated for thalamus of 22L-infected animals and normal controls for three time points.

In contrast to the Astrocyte Function or Microglia Activation pathways (Fig. 4), the Neuron and Neurotransmission pathway showed a very subtle response to prion infection (Fig. S7). Such subtle response could be, in part, due to a limited number of genes related to neurons and neurotransmission in the Neuroinflammation panel (80 genes), which may not capture neuronal dysfunctions to the full extent. Nevertheless, only a minor down- or up-regulation of individual neuronal function-related genes, of which many lack statistical significance, argue against a substantial loss of neuronal population even at the terminal stage of the disease.

To summarize, the Astrocyte Function pathway scored at the top of the list for both prion strains among the pathways analyzed by the Neuroinflammation panel. Moreover, changes in the genes associated with the Astrocyte Function pathways were detectable already at the preclinical stage of the disease. While Microglia Activation pathway scored very high too, its scores are likely to be inflated due to microglia proliferation.

### A1-, A2- and PAN-reactive markers are upregulated

Previous studies established that, depending on activation stimuli, astrocytes can acquire two opposite reactive phenotypes: proinflammatory, neurotoxic A1 state and neuroprotective A2 state [41, 42]. The concept of A1/A2 phenotypes has been employed for characterizing astrocyte activation states under pathological conditions or normal aging [7, 8]. We found that among the A1-, A2- and PAN-specific markers included in the Neuroinflammation panel, the majority of PAN-reactive markers (*Osmr, Vim, Serpina3n, Cxcl10, Timp1, S1pr3, Lcn2, Hspb1, Cp*) as well as several A1-specific (*Psmb8, Gbp2, H2-T23, Serping1*) and A2-specific (*Tgm1, Cd14, S100a10, Ptx3, Cd109*) markers were upregulated at the advanced stage of prion disease (Fig. 9). Both prion strains showed upregulation of the same markers suggesting that the reactive astrocyte phenotype lacked prion strain specificity (Fig. 9). However, the extent to which markers were upregulated in four brain regions mirrored the strain-specific dynamics of neuroinflammation: Th>HTh>Ctx>Hp for 22L, and Th> Ctx>Hp>HTh for ME7.

**Figure 9.**
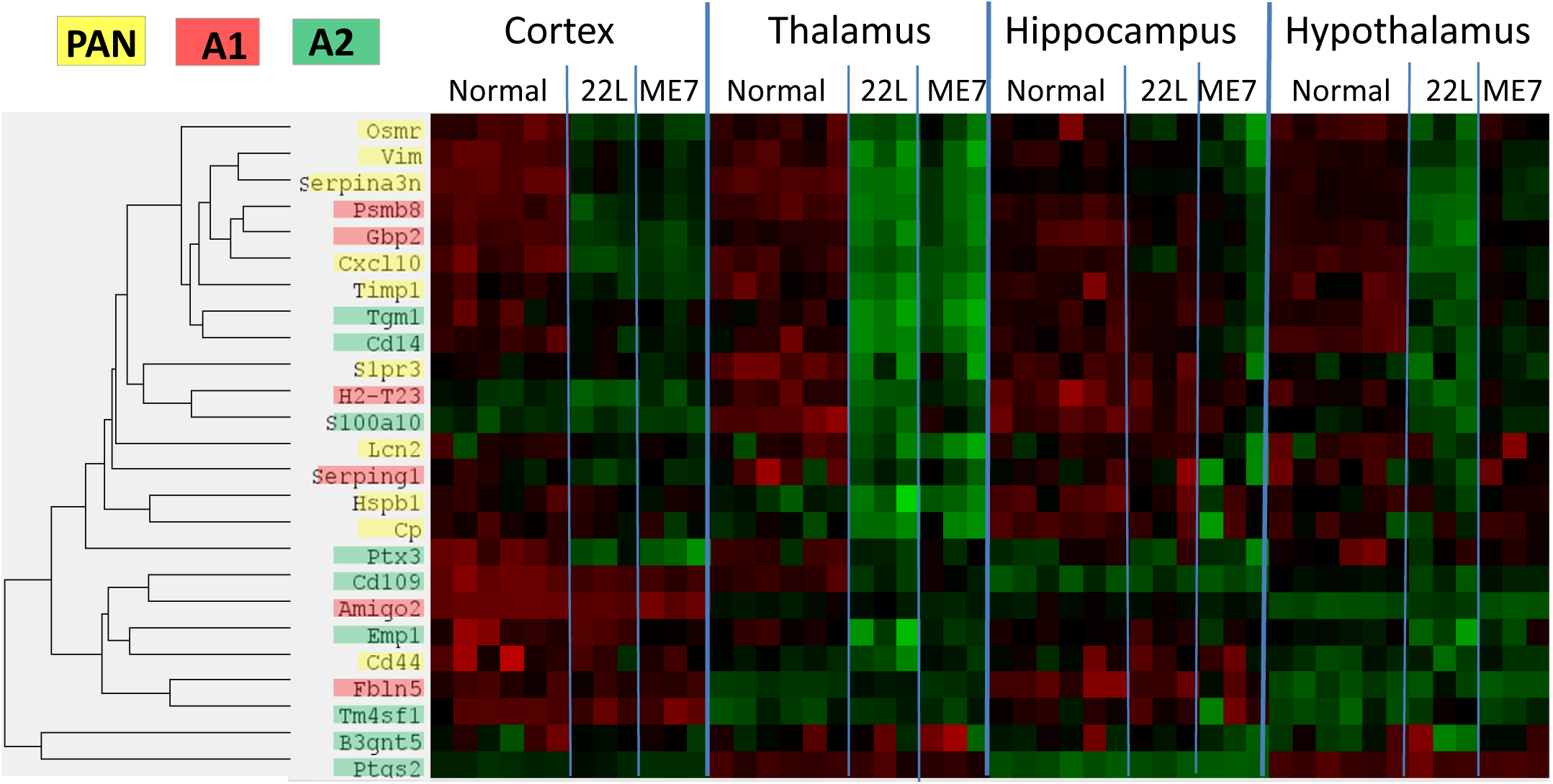
Heatmap of the A1-, A2- and PAN-specific markers. Heatmap analysis of A1-, A2- and PAN-specific markers in four brain regions of 22L-, ME7-infected animals and normal controls assessed for the third time point and shown for individual animals.

## Discussion

Recent advances in transcriptome analysis, single-cell RNA-sequencing and single-cell cytometry revealed considerable heterogeneity of glia phenotypes under normal conditions as well as dynamic changes in aging and neurodegenerative diseases. Single-cell transcriptional profiling of 1/2 million cells identified seven molecularly distinct and regionally restricted astrocyte types, in which regional specialization were found to be defined developmentally [9]. Single cell cytometry mapped distinct subsets of microglia populations providing insight into phenotypic heterogeneity among CNS-resident myeloid cell [43–45]. Moreover, genome-wide transcriptome analysis demonstrated that, like astrocytes, microglia too have distinct region-dependent homeostatic transcriptional identities [5]. Transcriptome profiling of human brains documented that, in astrocytes, the region-specific expression patterns undergo a major shift with normal aging [6]. Remarkably, region-specific differences in the expression of astrocyte-specific genes were found to largely disappear with old age [6]. Likewise, microglia isolated from mouse brains also showed diminishing of region-specific homeostatic signatures with normal aging [5]. Moreover, upregulation of microglia-specific genes and, in particular, those involved in immune and inflammatory functions (*C1q, Trem*) were found with age in humans [6]. Notably, a global shift in the expression pattern of glial-specific genes predicted age with greater precision than the expression of neuron-specific genes, underscoring the role of glia in normal aging [6]. Both microglia and astrocytes age in a regionally dependent manner showing variable aging rates in different regions [5, 8].

In the current study, we asked whether region-specific homeostatic identity of glial cells changes with progression of prion diseases, and if so, whether these changes occur in a region-dependent or uniform manner. Four brain regions of mice infected with 22L strain, which is mainly associated with astrocytes, or ME7, which is found in association with neurons [24], were examined at three time points. For both strains, all data sets collected for the first, preclinical time point separated into four distinct clusters in strict accordance to the brain regions, illustrating region-specific homeostatic signatures. However, appearance of the first clinical signs was accompanied by overexpression of a subset of genes forming neuroinflammation signature, which started to dominate over the region-specific homeostatic signatures. A departure from homeostatic region-specific identity strengthened further at the advanced stage of the disease. While manifested to a different extent, the same neuroinflammation signature was observed in all four regions examined. Moreover, both astrocyte-associated 22L and neuron-associated ME7 strains showed the same neuroinflammation signature suggesting that it is independent of strain-specific cell tropism. Nevertheless, while the neuroinflammation signature was region- and strain-independent, neuroinflammation spread across four brain regions in a strain-specific manner. In fact, at the advanced stage of the disease, the four brain regions were affected to a different extent in 22L- and ME7-infected animals showing strain-specific ranking order with respect to the severity of neuroinflammation. This study illustrates that, in a manner resembling normal aging, glia lose their region-specific identity with the progression of prion diseases, although, this process occurs at a much faster rate in animals infected with prions.

Comparison of the top differentially expressed genes from the current work (*Cxcl10, Serpina3n, Lag3*, *Fcgr2b, C1qa, C1qb, C1qc, C4a, Stat1*, *Trem2*, Table 1) with those in the previous studies showed excellent agreement [24, 27–31]. At the top of the list was *Cxcl10*, a proinflammatory chemokine that can contribute to neurotoxicity and apoptosis. *Cxcl10* was identified in previous studies that employed global gene expression approaches or targeted approaches [24, 31, 46]. *Serpina3n*, a member of a large family of serine protease inhibitors, is a part of astrocytic PAN-reactive gene panel, which is upregulated in normal aging [7, 8]. Mouse *Serpina3n* was activated in ME7-, RML- and 301V-infected mice [27, 31]. Its human homolog *Serpina3* was strongly upregulated at the mRNA and the protein levels in human prion diseases including variant CJD, sporadic CJD, iatrogenic CJD, familial CJD, Fatal Familial Insomnia and Gerstmann-Straussler-Scheinker syndrome, as wells as in BSE-infected macaques [47, 48]. Expression of *Lag3*, a lymphocyte activation protein 3 also known as CD223, was also found to increase in prion-infected brains, yet its knockout failed to modify disease progression [49]. *Fcgr2b* is involved in natural killer cell mediated neurotoxicity, and was previously shown to be upregulated in ME7- and RML-infected mice [27, 31]. *C1qa, C1qb* and *C1qc*, the subcomponents of the complement cascade factor *C1q*, and *C4a* component of the complement cascade were found to be upregulated in 22L-, RML-, ME7- and 301V-infected mice and 263K-infected hamsters [27, 31, 50, 51]. In periphery, PrP^Sc^ interaction with *C1q* is required for sequestration of prions by spleen and infection of follicular dendritic cells [52]. Deficiency of *C1q* delays the onset of the disease upon peripheral infection [53]. Upregulation of *Stat1*, a pro-inflammatory transcription factor involved in JAK-STAT pathway that mediates cellular response to cytokines and interleukins (including those produced in prion diseases), was found in 22L- or ME7-infected mice [54, 55]. *Trem2*, a triggering receptor expressed on myeloid cells-2, is a major genetic risk factor and a main player in Alzheimer’s disease (reviewed in [56]). Upregulation of *Trem2* was found in RML-inoculated mice [57]. Yet, its depletion, while attenuating markers of activated microglia, did not affect the incubation time or survival of prion-infected mice [57].

This study did not aim to identify new differentially expressed genes. However, comparison of the 22L thalamus at the third time point, the region with the most severe neuroinflammation, with the previous results on global gene expression in whole brain tissues [27, 30] identified 37 new upregulated (P<0.05, fold change >1.5) and 11 downregulated genes (P<0.05, fold change <0.66 fold) (Table S4). Improved sensitivity of detection could be due to a few factors. First, analysis of a brain region versus whole brain might improve detection of genes that are up- or down-regulated in a region-specific manner or genes that display significantly different levels of basal expression between different regions. Secondly, genes that respond regardless of a brain region but display statistically significant differential expression only in the most affected region might have a better chance of detection. Finally, Nanostring might provide a more sensitive way for detecting genes expressed at low levels than other approaches, as it directly counts the number of mRNA copies.

The prion-associated Neuroinflammation signature identified in the current work consists of genes that largely belong to the Inflammatory Signaling, Activated Microglia, Astrocyte Function and Autophagy pathways, but not the Neuron and Neurodegeneration pathway, which showed very modest response. Among neuron-specific genes, it is worth mentioning the upregulation of *Cidea* (Cell Death Inducing DFFA Like Effector A), which activates apoptosis, and downregulation of *Ngf* (Nerve Growth Factor) that helps neurons grow and survive. A modest scoring of the Neurons and Neurotransmission pathway was consistent with previous results on transcriptome analysis that revealed relatively few degenerating neurons even at the advanced stages of the disease [28]. The group of activated genes in the Inflammatory Signaling, Astrocyte Function, Activated Microglia and Autophagy pathways showed very similar expression dynamics in response to prion infection, as assessed in four brain regions monitored at three time points for two prion strains (Fig. 4). Such coherent dynamics is remarkable, as it suggests that the same mechanism is involved in responding to prion infection regardless of a brain region or a cell tropism of the prion strain.

A number of previous studies that employed animal models, post-mortem human brains or cells cultured *in vitro* aimed at defining role of microglia in prion diseases. Consistent with the current studies, activation and proliferation of microglia were found to mirror PrP^Sc^ accumulation with respect to the affected brain regions and timing of PrP^Sc^ accumulation [30, 37, 58–64]. Using purified, brain-derived PrP^Sc^, we previously showed that PrP^Sc^ can directly trigger proinflammatory response in primary microglia, and that the chemical nature of the carbohydrate groups on the N-linked glycans of PrP^Sc^ is important for microglia activation [38]. Nevertheless, the precise role of glia in chronic neurodegeneration associated with prion disease has been under extensive debate and remains controversial [39, 57, 65, 66]. Microglia activation was shown to occur at much earlier stages than synaptic loss [24, 34, 35, 37, 67], which is considered to be an early neuron-specific pathological sign [68, 69]. Solid evidence in support of both neuroprotective phenotypes and inflammatory or neurotoxic phenotypes have been presented over the years [58, 61, 63, 67, 70–78]. It is likely that multiple reactive phenotypes co-exist and undergo changes with disease progression. Several microglia transcripts upregulated in the prion-infected mice were shared with those in aged mice (*Irf7, Stat1, Ifitm* family) [5], or shared with aged mice together with the APP/PS1 model of Aβ amyloidosis and the AAV-Tau^P301L^ model of tauopathy (*Apoe, C3, Ccl3, Clec7a, Itgax, Lilrb4, Spp1)* [79]. Moreover, the neuroinflammation signature identified in prion-infected animals in the current study partially overlapped with the microglia degenerative phenotype (MGnD) and the disease-associated microglia phenotype (DAM) reported previously in mouse models of other neurodegenerative diseases [80–82]. All three microglia phenotypes showed upregulation of the following genes: *Apoe, Axl, Clec7a, Csf1, Fabp5, Grn, Lilrb4, Spp1, Trem2, Tyrobp* and downregulation of *Egr1.* Yet, significant differences were also found. Prion-infected animals upregulated the following genes that were downregulated in DAM and MGnD: *P2ry12, Ccr5, Csf1r, Cx3cr1, Gpr34, Tgfb1, Tgfbr1, Mertk, Tmem119, Sall1, Mafb*, and, vise versa, the gene *Vegfa* was downregulated in prion-infected animals while upregulated in DAM and MGnD [80–82]. Together, these results suggest that prion disease is characterized by prion-specific neuroinflammation signature, which only partially overlaps with those associated with other chronic neurodegeneration conditions.

Growing evidence suggests that autophagy plays a role in prion clearance on one hand, and regulates the release of prions via exosomes on the other [83–85]. Upregulation of the Autophagy pathway genes was found to occur in parallel with the genes of Activated Microglia and Astrocyte function pathways, suggesting that microglia and/or astrocytes are responsible for autophagosomal clearance. However, ontological analysis of genes in microglia isolated from prion-infected mice in a previous study did not identify autophagy among pathways activated in microglia [30]. Moreover, unlike astrocytes and neurons, microglial cells do not replicate prions and were found to exhibit very limited efficiency in phagocytosis of PrP^Sc^ questioning whether they are involved in PrP^Sc^ clearance [78]. Microglia serve as the main guard for protecting CNS against pathogens. To discriminate between mammalian host cells and invading pathogens, microglia employs the same strategy as macrophages that involves sensing sialylated and non-sialylated glycans on a cell surface [86–88]. Majority of pathogens can synthetize only non-sialylated glycans and, instead of sialic acid, expose galactose at the terminal positions of glycans, which triggers phagocytic “eat me” signal in microglia [89]. Like mammalian cells, PrP^Sc^ surface is heavily sialylated due to sialic acid residues at the terminal positions of the PrP N-linked glycans [21, 22, 90]. Previously, we showed that sialylation status of PrP^Sc^ is essential in determining the fate of prion infection in an organism [90–93]. Reducing of PrP^Sc^ sialylation was found to fully abolish its infectivity upon administration to animals [90, 92, 93]. It is not known, whether microglia can phagocyte PrP^Sc^ with normal sialylation status. Human astrocyte cell line was found to be capable of up-taking and degrading PrP^Sc^ [94]. Howevr, it is not know whether astrocytes senses the same cues as the cells of myeloid origin for activating their phagocytic machinery. It remains to be determined whether astrocytes play a major role in clearance of PrP^Sc^ *in vivo* and whether PrP^Sc^ clearance involves phagocytosis and autophagy.

In the current study, several components of a complement cascade including *C1qa, C1qb, C1qc, C4a* and *C3ar1* (receptor for C3a) were found to be strongly upregulated in prion-infected mice. Components of complement system including *C1q*, *C3* and *C4* play a critical role in synapse pruning during normal brain development [95, 96]. The developmental mechanisms of synaptic pruning involve tagging of synapses by C1q, their opsonization by C3, followed by their engulfment and phagocytosis via interaction with the C3 receptor expressed by microglia. Growing evidence suggest that similar mechanisms of synapse elimination are activated in reactive microglia in neurodegenerative diseases including Alzhemer’s diseases, frontotemporal dementia as well as normal aging [97–100]. Among other genes that might be involved in neurotoxic activity of microglia is *Axl*, a TAM tyrosine kinase receptor. Under normal conditions, *Axl* drives microglial phagocytosis of apoptotic cells during adult neurogenesis [101]. However, expression of Axl was found to be upregulated in mouse models of Parkinson’s disease [101] as well as in prion-infected mice in the current study. Upregulation of components of a complement cascade observed in the current work suggest that non-cell autonomous mechanisms might be responsible for synaptic loss and dysfunction in prion diseases.

Reactive astrogliosis, routinely observed as an increase in GFAP signal, is one of the central features of prion diseases, yet information about the role of astrocytes in prion disease is scarce. Astrocytes can replicate and accumulate PrP^Sc^ independently of neurons [102–105]. Furthermore, expression of PrP^C^ on astrocytes was found to be sufficient for prion-induced chronic neurodegeneration [106, 107]. However, it remained unclear whether normal physiological functions of astrocytes are altered as a result of reactive astrogliosis. Unexpectedly, the current study found the Astrocyte Function pathway scored at the top of the tested pathways. While 22L does replicate in astrocytes, ME7 is predominantly neurotropic. Therefore, change in the expression of genes associated with astrocyte function cannot be attributed directly to the replication of PrP^Sc^ by astrocytes. In homeostatic state, astrocytes are responsible for a variety of physiological functions including modulation of neurotransmission, formation and elimination of synapses, regulation of blood flow, supplying energy and providing metabolic support to neurons, maintaining the blood-brain-barrier, synthesis of cholesterol and much more [108, 109]. Because the representation of genes that report on astrocytic function in the Neuroinflammation panel is limited, the current work cannot identify specific functions that are disrupted or upregulated in the reactive astrocytes. We also do not know how changes in astrocyte function affects neurons.

Clustering analysis of the global scores at the first, preclinical point suggests that astrocytes might respond to prion infection ahead of microglia. While unexpected, this result is consistent with the previous findings that in mice infected with 22L, ME7 or RML, GFAP was upregulated prior of Gpr84, the marker of activated microglia [24]. While these findings need to be confirmed using more detailed analysis of the preclinical stage, this result suggests that the relationship between reactive microglia and astrocytes appears to be more complex than one could envision based on the Barres’s hypothesis. According to this hypothesis, microglial cells in proinflammatory reactive phenotype release astrocyte-activating signals (Il-1α, TNG-α and C1q), that induce pro-inflammatory A1 neurotoxic states in astrocytes [3, 41]. In support of this hypothesis, blocking of reactive microglia stimuli that induce A1 astrocytic phenotype was found to be neuroprotective in mouse models of Parkinson’s disease [110]. Moreover, consistent with this hypothesis, previous studies demonstrated attenuation of astrocytic gliosis and a delay of clinical onset of prion diseases in mice deficient of interleukin-1 receptor, which is activated by proinflammatory cytokines produced by microglia [111]. However, contrary to expectation, progression of prion disease was significantly accelerated in the triple (*Il1α^−/−^, TNFα^−/−^* and *C1q^−/−^*) knockout mice infected with prions [112]. This finding contradicts the Barres’s hypothesis and raises the question of whether astrocytes are followers or drivers of glia proinflammatory phenotype. The findings in the tripled knockout mice is unexpected, yet it could be reconciled with the Barres’s hypothesis, if one assumes that the type of reactive phenotype acquired by astrocytes in prion diseases upregulate phagocytosis, which might be important for prion clearance. Indeed, in brains subjected to ischemia, reactive astrocytes were found to upregulate genes responsible for phagocytosis and acquire phagocytic phenotype [113].

In a manner similar to M1 and M2 macrophage nomenclature, reactive phenotypes of astrocytes were formally classified as A1 (pro-inflammatory or toxic) or A2 (neurotrophic or neuroprotective) [3, 114]. In the current study, the majority of PAN-specific markers, as well as several A1 and A2-specific markers, were upregulated in both 22L- and ME7-infected groups. The extent of activation in four brain regions correlated well with the strain-specific degree of neuroinflammation in these regions. Analysis of A1-, A2- or PAN-specific markers in crude brain tissues should be considered with great caution, as most of the markers are not exclusively astrocyte-specific, but also expressed by other cell types. As such, these markers might be useful for reporting changes in region-specific microenvironments that govern astrocytic response under specific disease conditions. Nevertheless, the result of the current work is consistent with the recent study that also reported mixed astrocyte phenotype in prion disease [112]. It is not clear whether appearance of mixed phenotype, if such exist, could arise due to co-existing mixtures of pure A1 and A2 states, multiple distinct activation states in addition to classical A1 and A2 states, co-expression of markers of different phenotypes within individual cells, or all of the above. Observation of mixed astrocyte phenotype in prion disease is not entirely unexpected considering that the reactive phenotype of microglia only partially overlaps with those described in mouse models of other neurodegenerative diseases.

In future studies, it would be interesting to examine the extent to which neuroinflammation signature observed in prion infected animals resembles those in normal aging. Another area of considerable interest involves assessing changes in functional states of astrocytes and determining whether these changes are protective against neurodegeneration or contribute to neurodegeneration. The Neuroinflammation panel is dominated by the genes that report on microglial activation, so further analyses that focus on astrocyte function are warranted. Designing a panel with an emphasis on astrocyte-specific genes and/or analysis of acutely isolated astrocytes could shed a light on the role of astrocytes in chronic neurodegeneration. Defining causative relationship between reactive states of microglia and astrocytes is also of considerable interest.

## Conclusions

A growing number of studies demonstrated phenotypic heterogeneity in microglia and astrocytes under normal conditions as well as dynamic region- and subregion-specific changes in glia phenotypes under aging or neurodegenerative conditions. While region- and subregion-specific changes in glia phenotype has been documented using a number of animal models of neurodegenerative diseases, the extent to which animal models recapitulate neurodegenerative disease in human has been under intense debates. Among neurodegenerative diseases, prion disease is the only one that can be faithfully reproduced in wild type animals. Indeed, non-transgenic, inbreed mice infected with prions develop actual *bona fide* prion disease, and not a disease model. Yet, a significant gap in our understanding of glia phenotype in prion diseases exists. Previous transcriptome studies of prion-infected animals employed whole brain tissues for differential gene expression analysis, leaving region specific identities concealed. To fill the gap, the current study is the first to analyze temporal changed in neuroinflammation transcriptome in prion diseases in a region-specific manner. The current work revealed that (i) region-specific homeostatic identities of glia were preserved at the preclinical stages of prion disease. (ii) With the progression of clinical signs, region-specific signatures were lost and replaced with a uniform neuroinflammation signature. (iii) Neuroinflammation transcriptome signature was not only region-independent but also uniform for prion strains with different cell tropism. (iv) Changes in astrocyte function scored at the top of activated pathways. Moreover, astrocyte function pathway responded to prion infection prior to activated microglia. (v) Prion-associated neuroinflammation signature identified in the current study overlapped only partially with the microglia degenerative phenotype and the disease-associated microglia phenotype reported for animal models of other neurodegenerative diseases.

## Abbreviations

22L: Mouse adapted prion protein strain 22L
Ctx: Cortex
DE: Differentially expressed
GFAP: Glial fibrillary acidic protein
Hp: Hippocampus
HTh: Hypothalamus
ME7: Mouse adapted prion protein strain ME7
PrP^C^: Normal, cellular form of the prion protein
PrP^Sc^: transmissible, disease-associated form of the prion protein
PBS: Phosphate-Buffered Saline
Th: Thalamus

## Declarations

### Ethics approval and consent to participate

This study was carried out in strict accordance with the recommendations in the Guide for the Care and Use of Laboratory Animals of the National Institutes of Health. The animal protocol was approved by the Institutional Animal Care and Use Committee of the University of Maryland, Baltimore (Assurance Number A32000-01; Permit Number: 0215002).

### Consent for publication

Not applicable

### Availability of data and materials

All data generated or analyzed in this study are included in this published article [and its supplementary information files].

### Authors’ contributions

IB and NM designed the study and wrote the manuscript. KM and JC performed animal procedures and scored the disease signs. NM dissected animal brains. JC performed isolation of RNAs. NM and JC prepared brain slices, performed immunohistochemistry and Western blotting. NM analyzed the data. KM edited the manuscript. All authors read and approved the final manuscript.

## Acknowledgments

We thank Institute for Genome Sciences at the University of Maryland School of Medicine for processing samples using nCounter Nanostring platform.

## Supporting Information

**Table S1.**
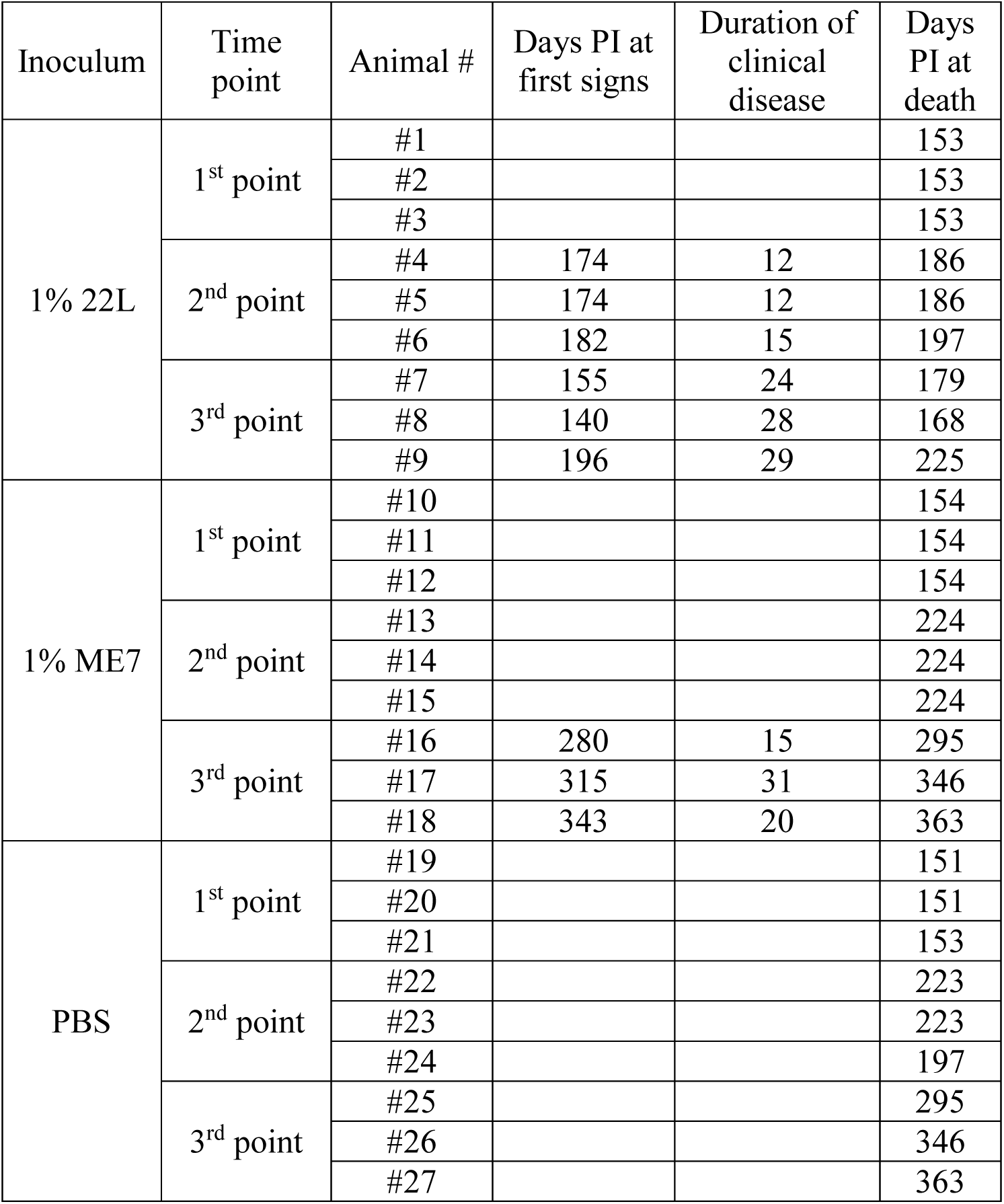
List of animals used for gene expression analysis.

**Table S2.** nCounter® Mouse Neuroinflammation Panel (in separate file)

**Table S3.** Top differentially expressed genes detected for the first time point (in separate file). Differentially expressed genes with P<0.05 are listed according to the prion strain and brain region.

**Table S4.** List of all differentially expressed genes in 22L- and ME7-infected animals in advanced stage of the disease (in separate file). Differentially expressed genes with P<0.05 are listed according to the prion strain and brain region. Newly identified differentially expressed genes (P<0.05, fold change >1.5 or <0.66) are highlighted in yellow boxes.

## Supporting Figures

**Figure S1.**
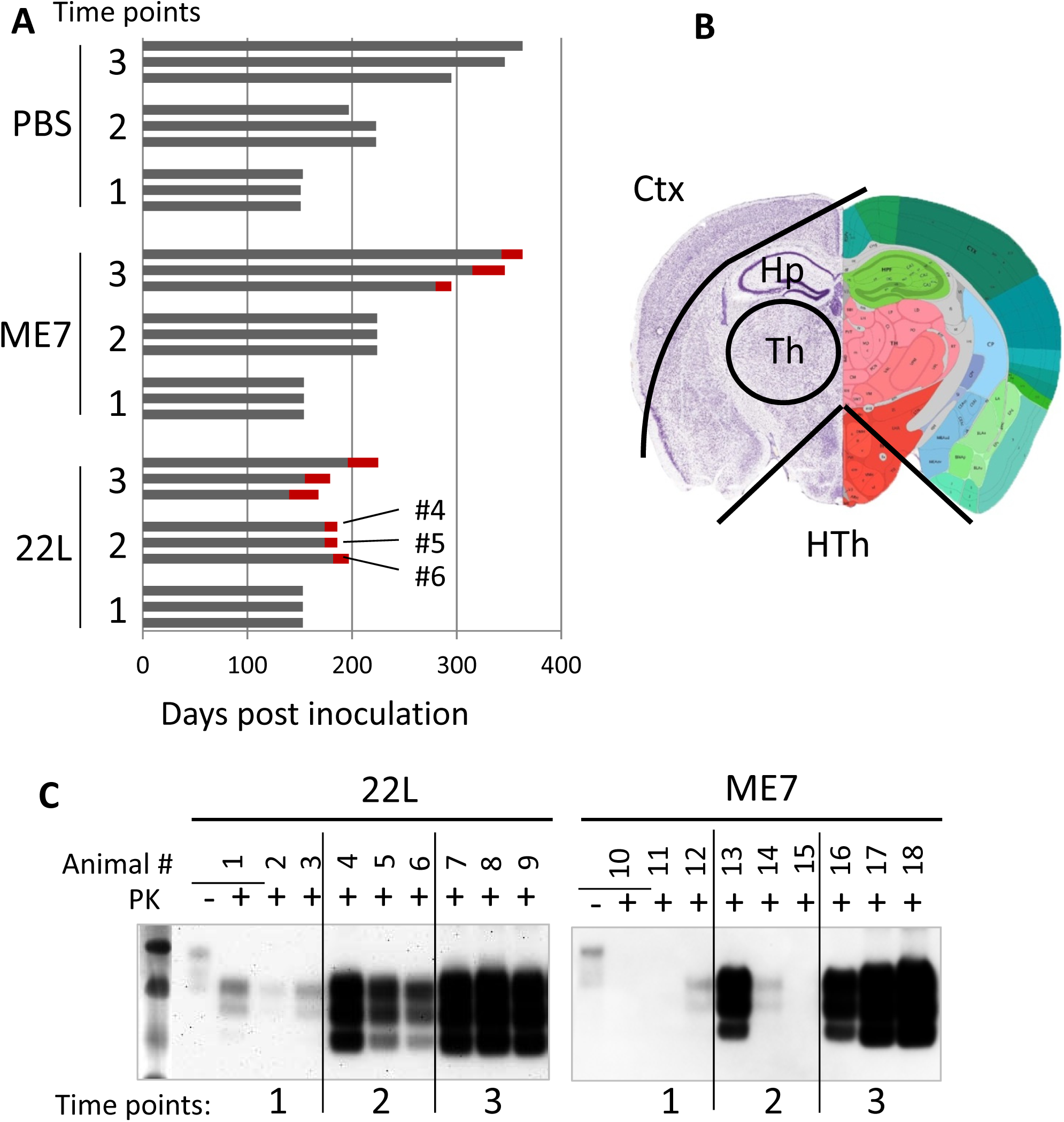
Experimental design. (**A**) Schematic representation of three animal groups inoculated IP with PBS, 22L or ME7 strains (200 μl, 1% brain homogenate) and euthanized at three time points post inoculation as indicated. (**B**) Coronal section of the brain showing four brain regions (Ctx - cortex, Hp - hippocampus, Th - thalamus, HTh - hypothalamus) collected for gene expression analysis. (**C**) Analysis of PrP^Sc^ amounts in animals used for the gene expression analysis. Western blots were stained with ab3531 anti-PrP antibody.

**Figure S2.**
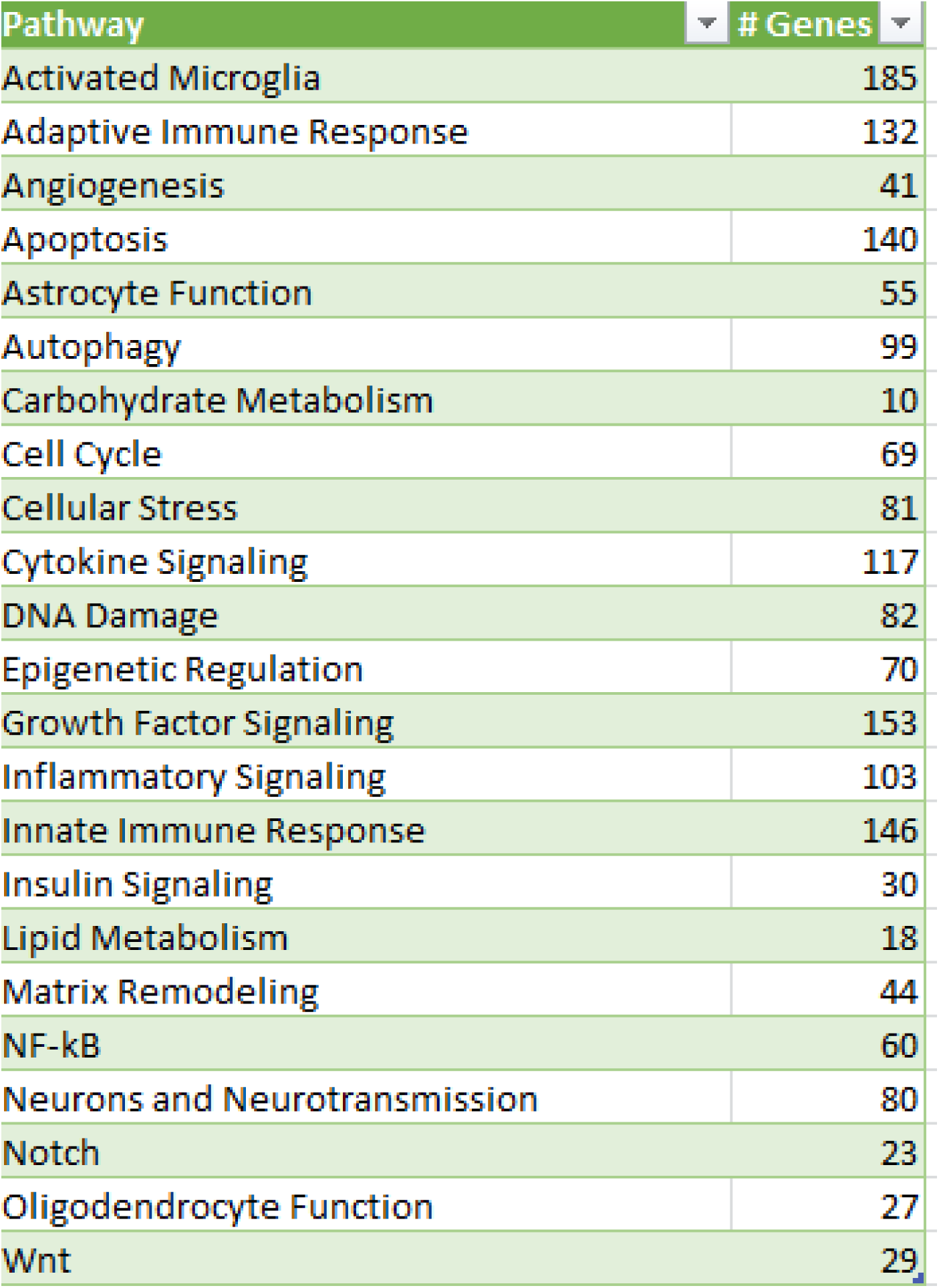
List of pathways with corresponding gene numbers analyzed by the Neuroinflammation panel.

**Figure S3.**
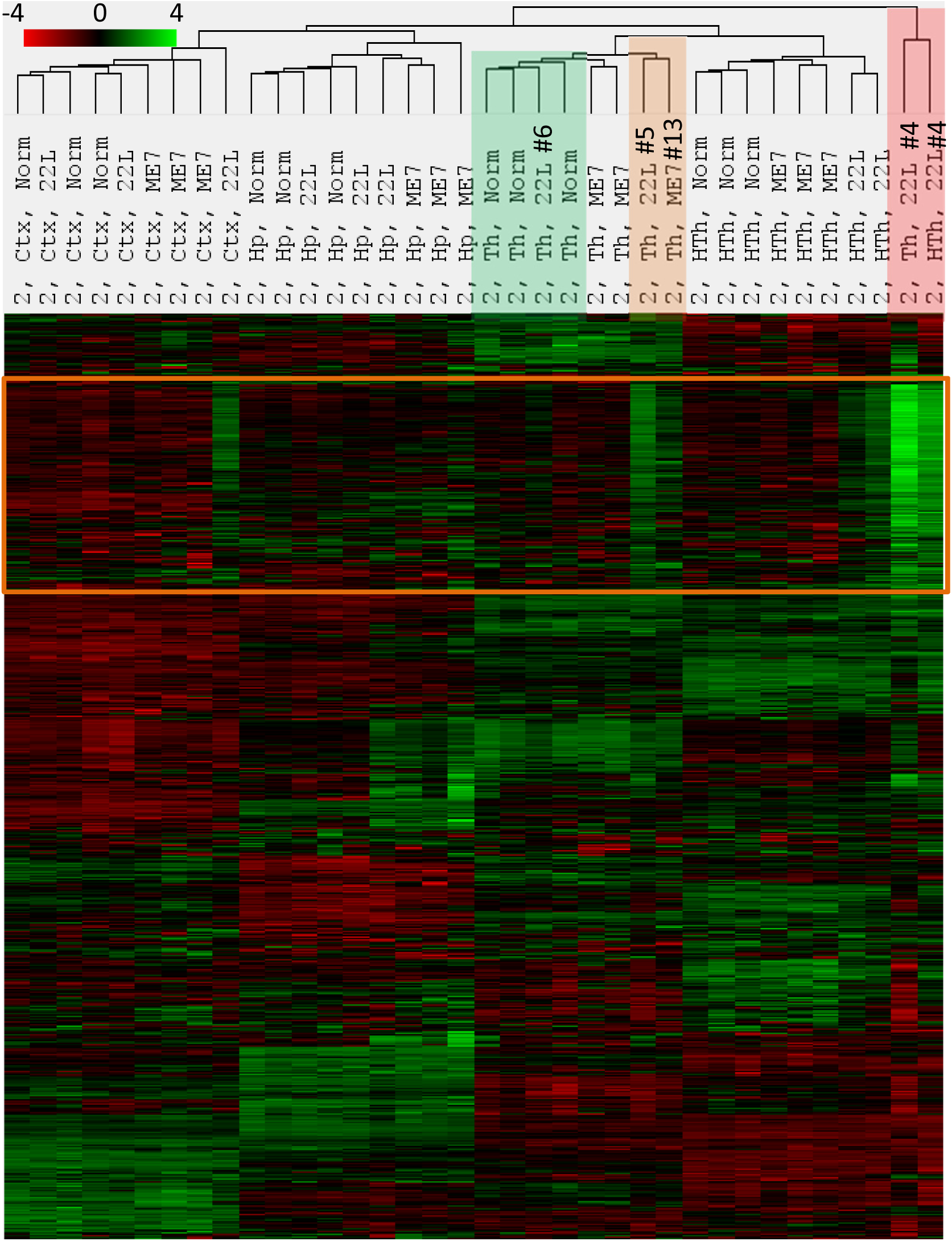
Agglomerative hierarchical cluster analysis of all data collected for the second time point. Animals from the second time point showed high variation in the degree of profile changes, which is particularly visible for a set of genes in the orange frame. Th and HTh from the most affected 22L-infected animal (#4) clustered away from other animals and brain regions (red shading), whereas Th from the least affected 22L-infected animal remained within the sub-cluster with normal Th. Th from 22L-infected, animal #5, and the most affected among ME7 group, animal #13, formed a separate sub-cluster within a Th cluster (brown shading).

**Figure S4.**
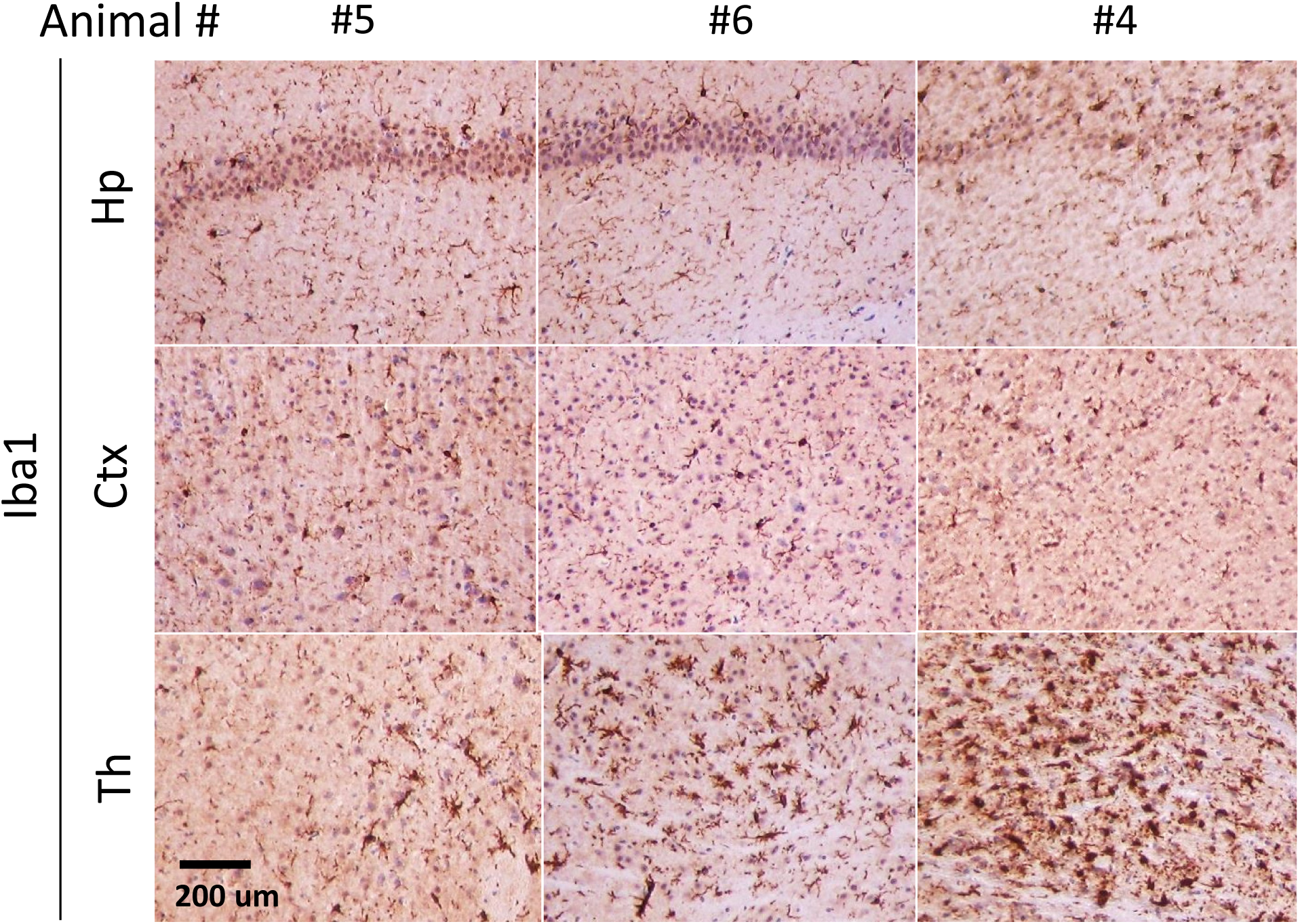
Histopathological analysis of 22L-infected animals from the second time point. Immunostaining of individual 22L-infected animals using antibody to microglia-specific marker, Iba1. Scale bars = 200 μm.

**Figure S5.**
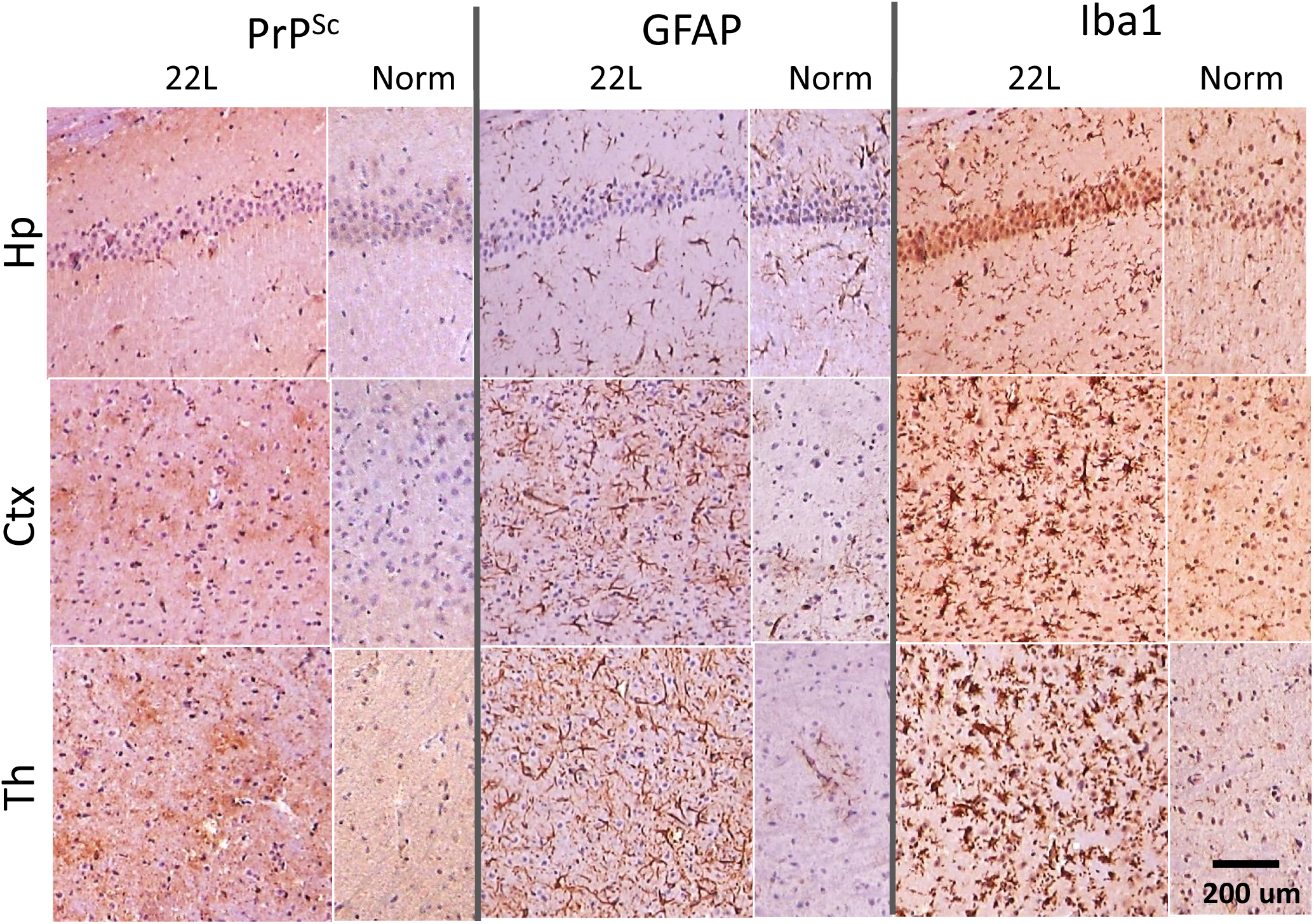
Histopathological analysis of 22L-infected animals from the third time point. Deposition of PrP^Sc^ probed by SAF-84 anti-PrP antibody, staining of microglia with anti-Iba1 or astrocytes with anti-GFAP in hippocampus, cortex and thalamus. Scale bars = 200 μm.

**Figure S6.**
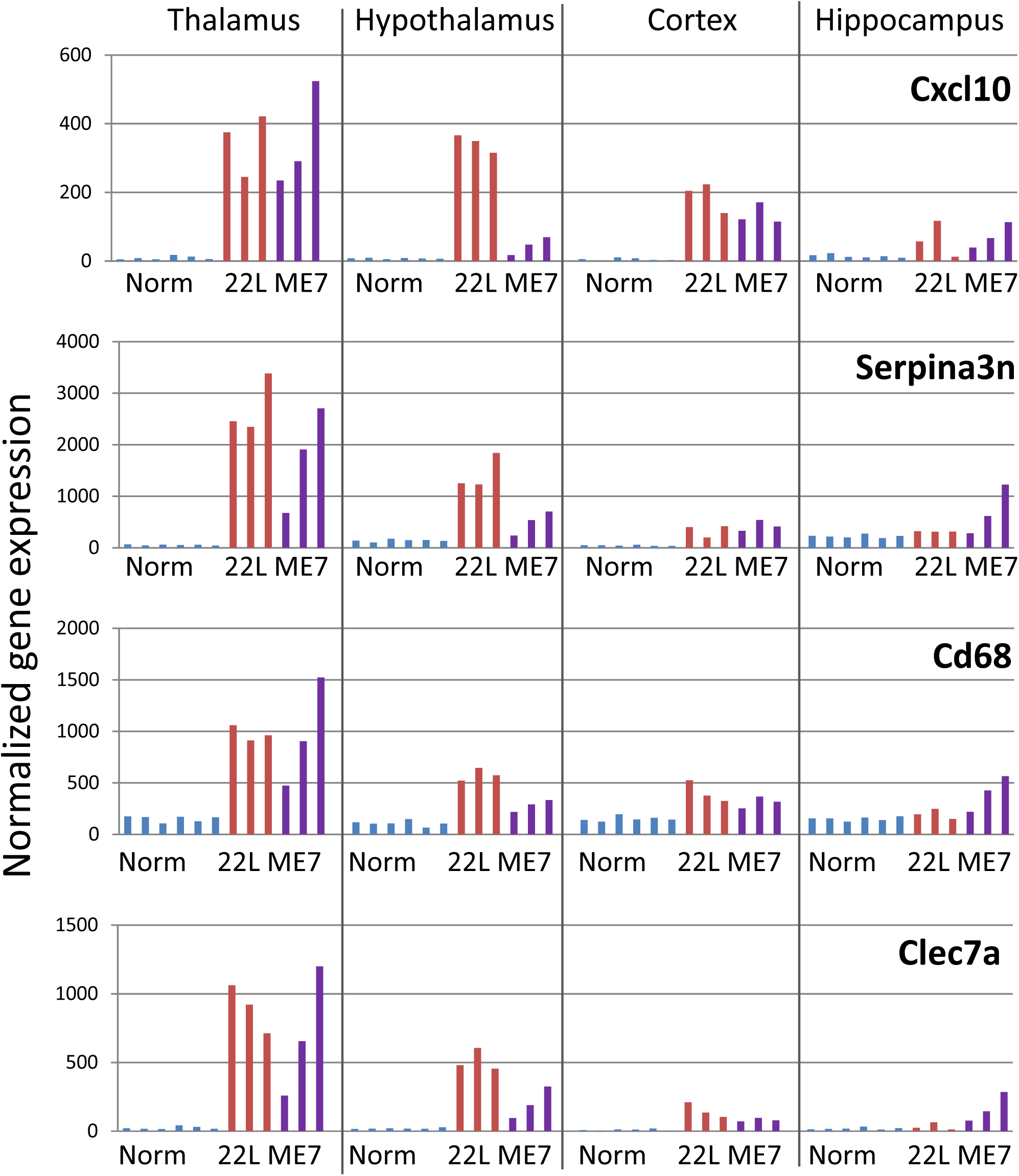
Region- and strain-specific pattern of expression of top differentially expressed genes. Normalized gene expression of *Cxcl10, Serpina3n, Cd68, Clec7a* in four brain regions in 22L- and ME7-infected animals and normal controls. Gene expression in individual animals is shown.

**Figure S7.**
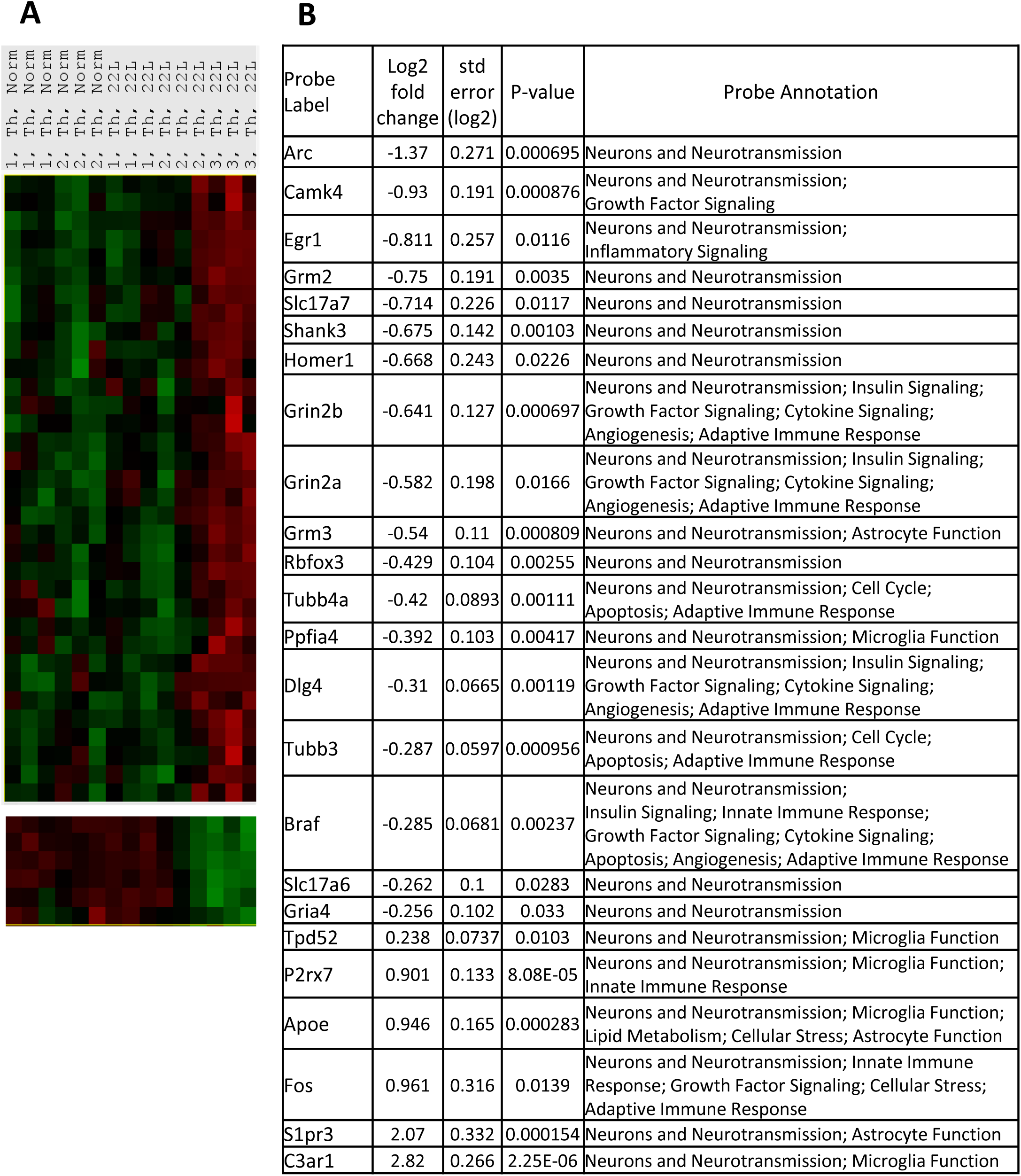
Differentially expressed genes in Neurons and Neurotransmission pathway. (**A**) Agglomerative clustering of the differentially expressed genes in Neurons and Neurotransmission pathway. Downregulated (top) and upregulated (bottom) clusters for thalamus at three time points are shown. (**B**) List of differentially expressed genes in Neurons and Neurotransmission with P<0.05 for thalamus at the third time point.

